# Pharmacologically induced weight loss is associated with distinct gut microbiome changes in obese rats

**DOI:** 10.1101/2021.05.06.442639

**Authors:** Silvia Raineri, Julia A. Sherriff, Kevin S.J. Thompson, Huw Jones, Paul T. Pfluger, Nicholas E. Ilott, Jane Mellor

## Abstract

**Background:** Obesity, metabolic disease and some psychiatric conditions are associated with changes to relative abundance of bacterial species and specific genes in the fecal microbiome. Little is known about the impact of pharmacologically induced weight loss on distinct gut microbiome species and their respective gene programs in obese individuals.

**Results:** Using shotgun metagenomics, the composition of the microbiome was obtained for two cohorts of obese female Wistar rats (n=10-12, total of 82) maintained on a high fat diet before and after a 42-day treatment with a panel of four investigatory or approved anti-obesity drugs (tacrolimus/FK506, bupropion, naltrexone and sibutramine), alone or in combination. We found that sibutramine treatment induced consistent weight loss through reducing food intake. Decreased weight loss in sibutramine-treated rats was associated with changes to the gut microbiome that included increased beta-diversity, increased Bacteroides/Firmicutes ratio and increased relative abundance of multiple *Bacteroides* species. In addition, the relative abundance of multiple genes was found to be differentially abundant, including significant reductions in components of flagellum and genes involved in flagellum assembly.

**Conclusions:** This study provides a large resource comprising complete shotgun metagenomics datasets of the fecal microbiome coupled with weight change and food intake at day 3, day 15 and day 42 from 82 obese rats treated with a range of compounds used for weight loss, which are available to the community for detailed analysis. Furthermore, by conducting a detailed analysis of the microbiome associated with sibutramine-induced weight loss, we have identified multiple weight-loss associated microbial taxa and pathways. These include a reduction in components of flagellum and the flagellum assembly pathway that points to a potential role of sibutramine-induced weight-loss on regulating bacterially driven anti-inflammatory responses.

## Introduction

The gut microbiome and human metabolism are tightly connected, as it is estimated that up to 10% of blood-circulating metabolites are products of reactions carried out by the microbiome [1]. Alterations to the microbiome have been linked to a wide variety of diseases, including obesity, inflammatory bowel disease and psychiatric conditions [2–4]. Whether disruption of the host microbiome is simply associated with a disease or is playing a causative role is not clear, although effort is currently focused on finding common patterns of microbiota alterations in individuals affected by a specific disease such as obesity, a pandemic affecting over 600 million adults worldwide [5][6][7–9]. The development of sequencing technologies such as 16S and shotgun metagenomics have allowed for descriptive analyses of the differences between the microbiota of obese and lean individuals, although progress is limited by high individual variability, and a lack of guidelines for study design and bioinformatics analysis. Nevertheless, a few common patterns have emerged. Obese patients show a less diverse microbiome, as both their α diversity, the number of different microbial species found within a subject, and their β diversity, the degree of species diversity between individuals, is decreased compared to healthy controls [3]. Obesity is associated with higher ratio of Firmicutes to Bacteroidetes phyla, although some studies reported little or no change [10–13]. Microbes from the Firmicutes phylum are more efficient at digesting dietary polysaccharides and crosstalk with the host, promoting the increased storage of triglycerides within adipocytes leading to obesity [14,15]. Finally, an increased presence of members of the Proteobacteria phylum, such as *Desulfovibrionaceae* and *Enterobacteriaceae*, may contribute to the obesity-induced inflammation state in patients. These Gram-negative bacteria add to the host microbiome a panel of genes associated to lipopolysaccharide (LPS) and peptidoglycan biosynthesis. A release of LPS into the bloodstream would elicit activation of Toll-like Receptor 4 (TLR4), thus igniting an inflammatory response [16,17]. Furthermore, fecal microbiota transplantation (FMT) studies provide evidence of a link between obesity and an altered microbiome [18].

Many studies have focused on describing the changes in the microbiome of obese patients after surgical procedures, such as bariatric surgery or Roux-en-Y gastric bypass, but less is known about the consequences of pharmacological treatment (Table S1) [11,17,19]. One therapeutic option is based on the combined treatment with bupropion and naltrexone, a dopamine uptake inhibitor and opioid receptor antagonist, respectively, both acting through suppression of appetite [6,20]. There is also anecdotal evidence for other compounds such as Tacrolimus, formally known as and from here onwards referred to as FK506, in weight loss [21,22]. High doses of FK506 act as an immunosuppressant, but its repeated administration at lower doses induces weight loss in rats (Supplementary Figure S1).

Here, we use shotgun metagenomics to sequence the microbiome of two cohorts of dietary-induced obese female Wistar rats, a rigorous animal model for obesity, before and after a 42-day treatment with a panel of drugs associated with weight loss to assess any changes induced in the microbiome [23]. Bupropion and naltrexone have been approved by the FDA and are currently used in the clinic as a treatment for obesity [8]. Sibutramine is an oral anorexiant and while still available in some countries, in many others it has been withdrawn from the market following its association with adverse effects [20]. FK506 was examined for its potential re-purposing as a weight loss drug [21,23]. These drugs are mechanistically associated with immunosuppression, amelioration of alcohol, opiate or nicotine dependence, and treatment of major depressive disorders (Table S1). We also assessed the effect of treatment with our panel of weight-loss associated drugs on the weight, food intake and glucose tolerance in obese rats maintained on a high fat diet, so that any changes to the microbiome could be related to physiology. Choosing shotgun metagenomics, a technique that allows the sequencing of all the genes in the fecal sample, as opposed to amplifying only the 16s rRNAs, allowed us to identify not only the species composition for each sample, but the individual genes as well, adding information regarding the functional potential across rats and between different treatment conditions. We present a large metagenomics dataset that will inform direct and indirect effects of these treatments on the composition of the gut microbiome and the functional role of each species.

## Materials and Methods

### Fecal collection protocol

Fecal samples from 82 female Wistar rats were collected at three timepoints during the in vivo experiment. On the mornings of Days −2, 15 and 42 immediately prior to dosing, a fresh cage pad was placed on the tray beneath the wire grid floor of the cage. The following morning (i.e. 24 hours later) approximately 15 faecal pellets were removed, placed into a sterile 50 ml tube and stored at − 80°C until DNA extraction was performed.

### In vivo experimental design

The in vivo experiments were carried out by RenaSci (Nottingham, United Kingdom) on behalf of Chronos Therapeutics Ltd. In order to induce obesity, animals had free access to powdered high fat diet (VRF1 plus 20% lard), ground chocolate, ground peanuts and tap water at all times. One hundred and six female Wistar rats (weight range 250-300 g; Charles River, Margate, Kent) were housed in pairs for 17 weeks for the induction of obesity. Eighty-seven animals were then selected for further experiments and housed singly in polypropylene cages with wire grid floors to enable the food intake of each rat to be recorded. After a two-week acclimation period, animals were divided into eight treatment groups and treated for forty-two days (Table 1). The panel of weight-loss drugs used includes FK506, bupropion, naltrexone and sibutramine, either alone or in combination.

**Table 1.** Treatment groups. Dosages and routes for drug administration of sibutramine and FK506 were based upon previous studies (Supplementary Fig. 1), while bupropion and naltrexone dosages were adapted from literature [24,25]

| Group | Treatment 1 (4 ml/kg po) | Treatment 2 (2 ml/kg ip) | Ref | n |
| --- | --- | --- | --- | --- |
| A | vehicle (castor oil/dehydrated alcohol po) | vehicle (saline ip) (S1) | S1 | 12 |
| B | FK506 (0.6 mg/kg po) | vehicle (saline ip) (S1) | S1 | 10 |
| C | vehicle (castor oil/dehydrated alcohol po) | bupropion (20 mg/kg ip) | [24] | 10 |
| D | vehicle (castor oil/dehydrated alcohol po) | naltrexone (1 mg/kg ip) | [24] | 10 |
| E | FK506 (0.6 mg/kg po) | bupropion (20 mg/kg ip) | S1[24] | 10 |
| F | FK506 (0.6 mg/kg po) | naltrexone (1 mg/kg ip) | S1[24] | 10 |
| G | vehicle (castor oil/dehydrated alcohol po) | bupropion (20 mg/kg ip) plus naltrexone (1 mg/kg ip) | [24] | 10 |
| H | sibutramine (5 mg/kg po) | vehicle (saline ip) | S1 | 10 |

Overall food intake in response to the drug treatments was calculated as the difference in weight of the chocolate, peanuts and High Fat Diet (HFD) pots in the rats’ cages between a certain day and the day before, then multiplied by the food energy values 20.79 kJ/g (HFD), 23.44 kJ/g (chocolate) and 30.34 kJ/g (ground peanuts). The sum of these three values is the combined food intake for a particular day. Throughout the duration of the study, animals were maintained on a reverse phase light-dark cycle (lights off for 8 h from 09.30-17.30 h) during which time the room was illuminated by red light.

### Oral glucose tolerance test (oGTT)

An oral glucose tolerance test (oGTT) was performed on Day 37 for each cohort using a random selection of rats per group (n = 10-12, total of 81 animals per cohort in oGTT). On Day 36, rats were deprived of food overnight before the oGTT but allowed free access to water. The following day (Day 37), the 41 fasted animals were subject to the oGTT. Animals were cannulated and a first blood sample was taken (Timepoint −60). Animals were then dosed with vehicle or drug as appropriate (60 min before the glucose challenge). Four minutes before the glucose challenge, an additional blood sample was taken (Timepoint 0). Thus, baseline blood samples were taken both before vehicle/drug treatment and before the glucose challenge at 0 min. Animals were then dosed orally with D-glucose (2 g/kg) and further blood samples were taken 10, 20, 30, 45, 60 and 120 min later. Between blood-sampling animals were returned to the home cage with free access to water but not food. At all time-points, blood samples (~90 μl) were collected, and plasma separated by centrifugation (2,400 g for 5 min at 4°C) to produce a single aliquot of plasma (~45 μl) which was frozen (−80°C) and subsequently assayed for glucose and insulin as described below. After the final tail vein bleed, the cannulae were removed and feeding jars were returned to the animals. All cages were then moved back to the holding room (reverse-phase lighting). On the day of the oGTT, the other cohort of rats was dosed as normal.

### Measurement of Plasma Glucose and Insulin

Plasma glucose was determined using a clinical glucose assay reagent (Thermo Electron Infinity stable reagent TR15421, Thermofisher) in a 96-well format, performed according to the manufacturer’s instructions. Plasma samples were diluted in saline prior to analysis (10x) and all determinations were performed in duplicate. Optical density at 340 nm and 400 nm (correcting wavelength) was determined using a SpectraMax 340PC384 microplate reader (Molecular Devices, USA). The optical density change was then calculated by subtraction.

Insulin was determined using the ultrasensitive rat insulin ELISA kit (cat. No 10-1251-01, Mercodia, Sweden) according to the manufacturer’s instructions, with additional insulin standards (1.50 and 1.75 ng/ml) to extend the range of the assay. Plasma samples were diluted 10x in saline solution prior to analysis, then assayed as single replicates, whilst standards were in duplicate. A five-parameter logistic curve of optical density against concentration was used to create standard curves using statistical analysis software (SAS®), then sample concentrations were estimated from the curves.

### Fecal DNA extraction

Fecal DNA was extracted from randomized samples using the QIAmp PowerFecal DNA kit (Qiagen, Hilden, Germany), following the manufacturer’s instructions. DNA was quantified on a Nanodrop (Thermofisher, Massachusetts, USA), then diluted to a final concentration of 10 ng/ μl. To get 200 base-pairs (bp) fragments, samples were sonicated (Bioruptor, Diagenode, Belgium) with the following settings: 70 mins, 30” on/30” off, Low followed by 20 mins, 30” on/30” off, Medium. DNA was purified using a QIAQuick PCR Clean up and Purification kit (Qiagen) and quantified using Qubit (Thermofisher).

### Shotgun metagenomics sequencing

Library preparation was performed with 150 ng of starting material using NEBNext Fast DNA Library Prep Set for Ion Torrent (New England Biolabs, Massachusetts, USA) and following the manufacturer’s instructions. A barcode adapter (Ion Xpress Barcode Adapters 1-16, Thermofisher) was ligated to each library to allow for multiplexing. DNA fragments of around 250-300 bp were size-selected on an E-Gel™ SizeSelect™ II Agarose Gels 2% (Thermofisher) as an intermediate step of the library preparation. Final concentration of each single-end library was assessed on a High Sensitivity DNA Chip for Bioanalyser (Agilent, California, USA). Shotgun metagenomics sequencing was performed on an Ion Proton sequencer (Ion Torrent, Thermofisher). Eight to ten DNA libraries with different barcodes were pooled together at a final concentration of 250 pM. Pooled libraries were loaded on an Ion PI v3 sequencing chip using the Ion Chef System (Thermofisher). A single sequencing run was performed for each pooled library.

**Table 2.** Average weight before (Day -3) or at the end of the experiment (Day 42). Table describing the starting or final average weight for each treatment group. A third column shows P.values of the ANOVA statistical test (weight ~ Timepoint + Cohort).

| Treatment Group | Average Weight<br>(Day -3) | Average Weight<br>(Day 42) | ANOVA adjusted<br>P.value |
| --- | --- | --- | --- |
| Ctrl | 420.30 | 435.69 | 0.338 |
| FK506 | 420.6 | 416.27 | 0.813 |
| bupropion | 417.45 | 431.56 | 0.512 |
| naltrexone | 416.38 | 436.76 | 0.309 |
| FK506 & bupropion | 421.61 | 408.54 | 0.443 |
| FK506 & naltrexone | 420.05 | 428.79 | 0.665 |
| bupropion & naltrexone | 421.03 | 433.39 | 0.533 |
| sibutramine | 420.12 | 391.83 | 0.0954 |

**Table 3.**
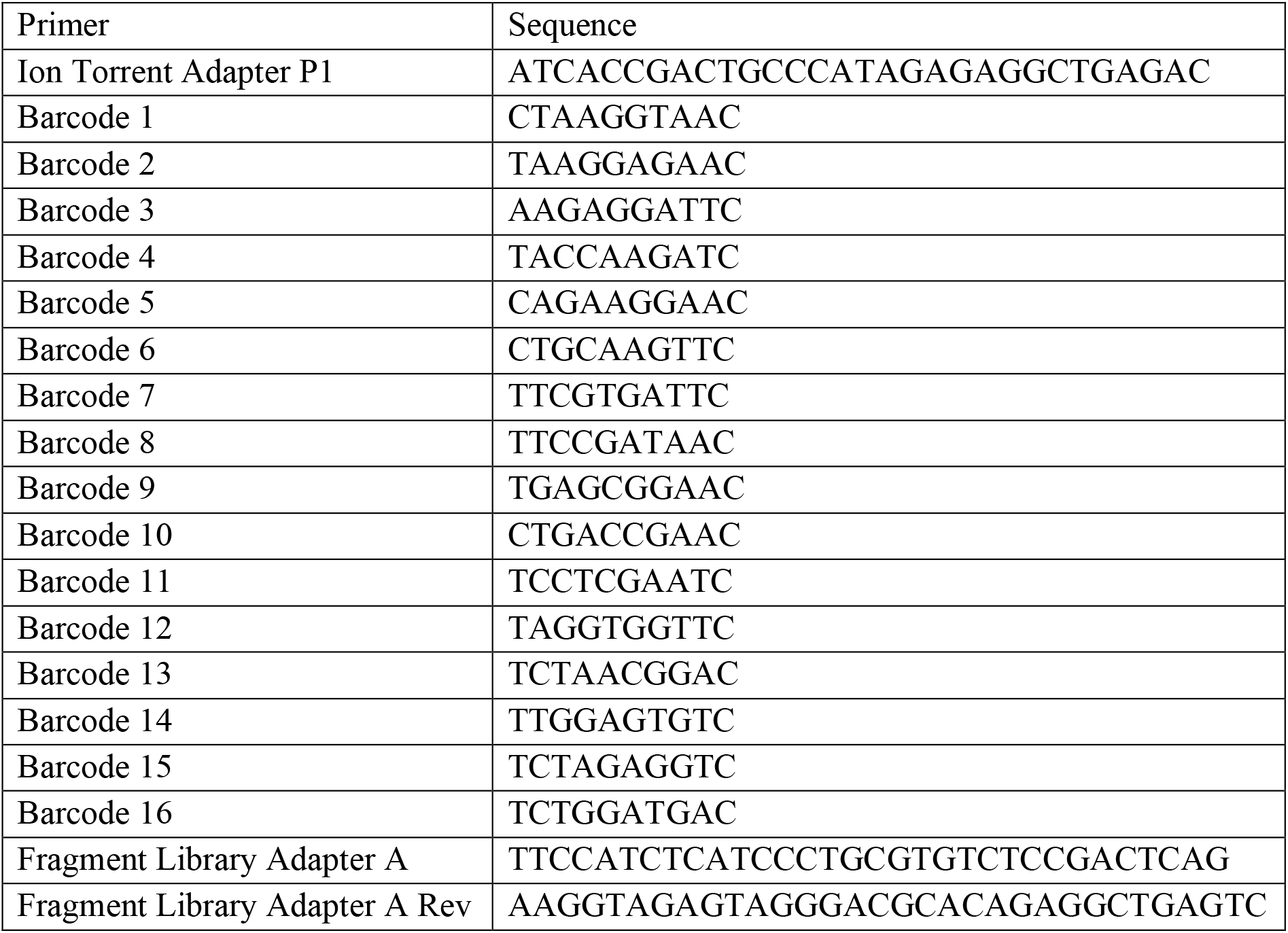
Primers and Barcodes used for library preparation and sequencing.

**Table 3.** ANOVA results for combined food intake consumption. Statistical significance for differences in food intake was computed with ANOVA using the model: value ~ Day + Treatment + Day *Treatment + sample within the Rstatix R/Bioconductor package. Resulting p-values were corrected into adjustedp-values (p.adj) using the Benjamini-Hochberg correction. Adjusted P-values are shown for every comparison in the model, together with generalized effect sizes (GES), representing the proportion of variance explained.

| ANOVA | Day |  | Treatment |  | Day by Treatment |  |
| --- | --- | --- | --- | --- | --- | --- |
|  | P.adj | GES | P.adj | GES | P.adj | GES |
| FK506 | 2.33E-21 | 0.18 | 3.40E-05 | 0.01 | 1.75E-02 | 0.07 |
| bupropion | 5.50E-20 | 0.18 | 0.21 | 0.002 | 0.88 | 0.036 |
| naltrexone | 7.84E-24 | 0.2 | 0.35 | 0.001 | 0.61 | 0.045 |
| FK506 & bupropion | 2.11E-21 | 0.19 | 0.003 | 0.01 | 0.24 | 0.05 |
| FK506 & naltrexone | 1.18E-16 | 0.16 | 0.48 | 0.0009 | 0.65 | 0.044 |
| bupropion & naltrexone | 6.63E-23 | 0.19 | 0.65 | 0.0002 | 0.65 | 0.04 |
| sibutramine | 1.16E-42 | 0.28 | 4.06E-27 | 0.11 | 6.56E-23 | 0.19 |

### Bioinformatics Analysis

#### Preprocessing files

Sequencing fastq files were loaded on Galaxy [26] (https://usegalaxy.org) for quality control. FASTQC (Galaxy Version 0.72) [27] was used to assess sequencing quality and evaluate the presence of adapters or other contaminating sequences. Sequences were trimmed using Trimmomatic (galaxy version 0.38.0) [28], using the SLIDINGWINDOW and MINLEN features to discard sequences below 30 bp long or whose average quality across a 4-nucleotide window drops below a defined threshold of 20. Newly trimmed sequences were then re-run on FASTQC to assess final quality. Host-derived sequences were removed from files through alignment with Bowtie2 (Galaxy Version 2.3.4.1) [29] against the rat canonical genome (Rn4). Unaligned sequences were written in a separate fasta file, which has been used as input for further analyses.

#### Taxonomic and functional profiling

Reads were processed using a cgat-core based pipeline (https://github.com/microbialman/MetaSequencing) to assemble reads into contigs, annotate those contigs with taxonomic and functional information and produce counts tables for each category of annotation [30]. Within this framework, reads were assembled into contigs using megahit with default parameters (v1.1.3) [31], which produced 44,789 (± 29,626) contigs of greater than 200bp in length. Open reading frames were identified in contigs using prodigal (v2.6.3) [32] with default parameters. Open reading frames were annotated with functional information using eggnog mapper (emapper v2.0.0) [33]. The Kyoto Encyclopedia of Genes and Genomes (KEGG) was used for downstream analyses. Contigs were annotated with taxonomic information using kraken2 [35] against the kraken2-microbial-fatfree database available from https://lomanlab.github.io/mockcommunity/mc_databases.html that contains “representative” or “complete” genomes across bacteria, archaea, fungi, protozoa, virus and UniVec_Core (i.e. potential vector contaminants in genome assemblies) domains.

Reads from each sample were aligned to the corresponding assembled contigs using bowtie2 (v2.3.4.1 with default parameters) [36], and counts tables were produced using eggNOG mapper- and kraken2-derived features (KEGG orthologous groups and taxonomy) using featureCounts (v1.6.0 with default parameters, [37]). Counts across samples were combined based on feature identifiers with a sample given a count of zero if the feature was not present. The resulting tables contained 6391 unique KEGG ko features, 54 phyla, 122 classes, 254 orders, 559 families, 1621 genera and 4850 species.

#### Microbiome composition analysis

Taxonomic profiling resulted in a table collecting microbiome composition for each sample at the species level. Diversity analyses were carried out using the R/Bioconductor package Phyloseq [38]. α diversity was calculated using Simpson’s diversity index through the estimate_richness function within the Phyloseq package, while β diversity was assessed using Bray-Curtis dissimilarity index to obtain a distance matrix (vegan, R package) [39], then visualized in a two dimensional ordination (PCA, base R). As sibutramine treatment displayed the greatest effect on weight loss, this condition was prioritized for downstream microbiome analysis. Differentially abundant genes or enriched species between sibutramine-treated rats at Day −3 compared to Day 42 were assessed using DESeq2 [40], with the following linear model: ~ rat id + Day. Heatmaps representing differentially abundant genes or enriched species were produced with the Heatmap3 Bioconductor package [41]. Analysis of species contribution to differentially abundant genes of interest were performed using a genes by species table obtained after taxonomic profiling, then absolute raw counts were transformed into relative counts. The top 10 contributing species were chosen for further representation, while the remainder was collapsed into the “other” category. KEGGREST Bioconductor package was used to decode KEGG-based KO gene identifiers into each gene name [42]. Statistical analysis was conducted using a combination of the stats, ggpubr and tidyverse R packages [43,44].

### Availability of data and materials

Raw fastq files are available from the ENA database, under the accession number PRJEB40767.

All scripts are available from a dedicated github page at the URL: https://github.com/SilviaRaineri/Microbiome_PRJEB40767. Processed data is uploaded on figshare and linked to the same github page.

## Results

### Only sibutramine induces significant weight loss in obese female rats maintained on an HFD

Initially, we monitored weight loss in two successive cohorts of female obese Wistar rats treated with weight loss drugs (Table 1). Routes and doses for the respective drugs were based on published pre-clinical studies in rodents/rats (Supplementary Figure S1)[23]. When considering both cohorts together, only sibutramine induced a statistically significant weight loss by the end of the study at Day 42, while rats in the Control or naltrexone groups significantly gained weight by Day 42 (Fig. 1).

**Figure 1.**
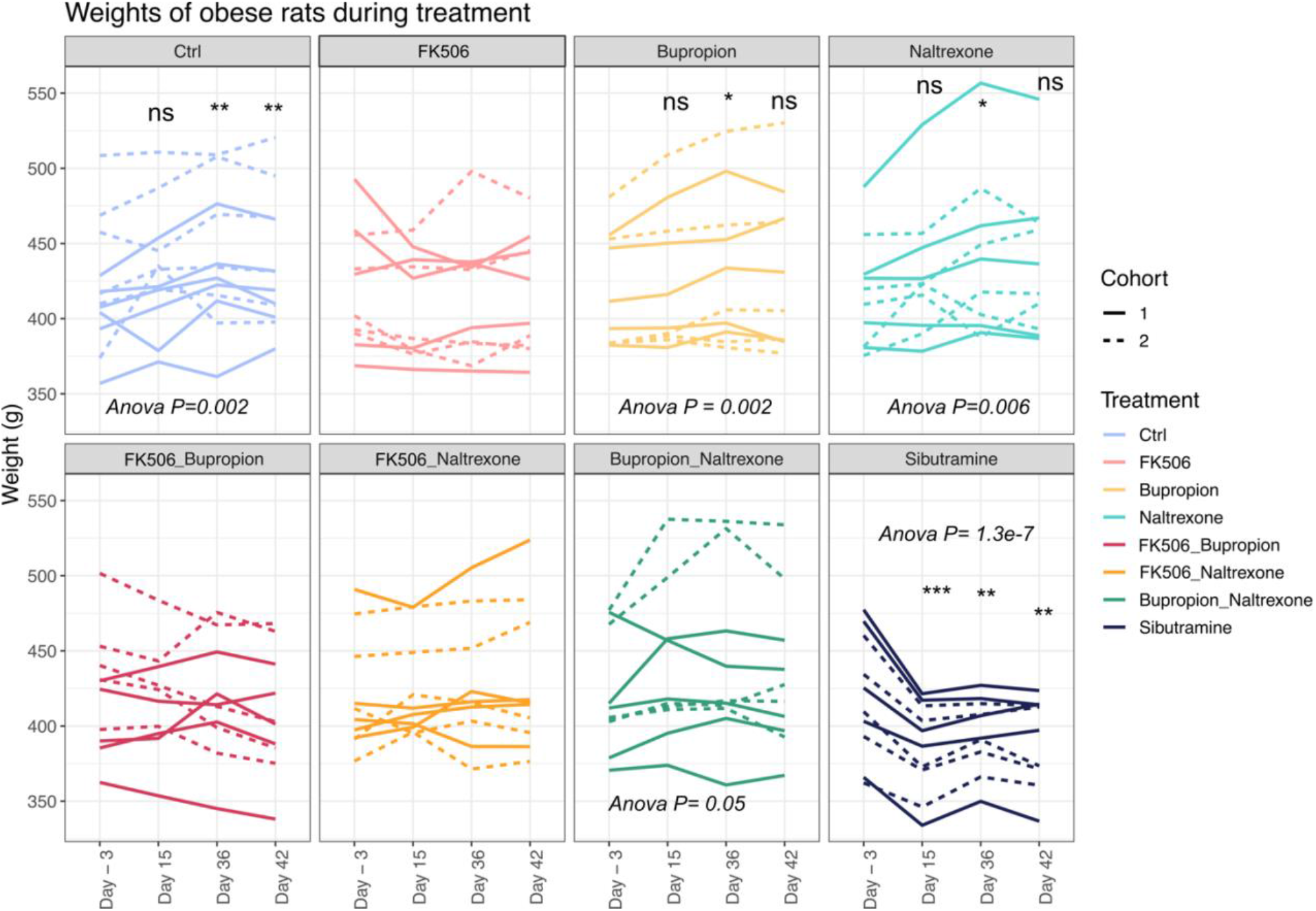
Weights of rats treated with anti-obesity drugs. Each panel shows the weights of rats treated with the respective weight loss drug at specific time points over the course of the study, starting three days before the first administration of the drug, then at Day 15, 36 and 42, the final day of the treatment. Statistical significance has been assessed first by repeated measures Anova, using the formula: weight ~ Timepoint + Rat number + Cohort, then significant timepoints were calculated with a series of paired t-tests, comparing Day – 3 weights with those at Day 15, 36 or 42. Paired T-test analysis within rats from the same treatment group showed only sibutramine treatment significantly reduced weight by the end of the treatment (Day -3 vs Day 42,Benjamini-Hochberg corrected p.values: Ctrl = 0.005 (T-statistic = −4.31, final weight significantly higher), FK506 = 0.52, bupropion = 0.06, naltrexone = 0.04 (T-statistic = −2.9, final weight significantly higher), FK506 and bupropion = 0.14, FK506 and naltrexone = 0.17, bupropion and naltrexone = 0.22, sibutramine = 0.005(T-statistic = 4.06, final weight significantly lower). Statistical significance levels: ns P > 0.05; * P < 0.05; ** P < 0.01; *** P < 0.00 (Control: N=12; sibutramine: N=10 rats per group).

Consistent with earlier reports showing a temporary weight lowering effect of sibutramine [45], we found a significant decrease of body weight within the first 15 days that was maintained throughout the duration of the study (ANOVA pvalue = 1.3e-7, Table 2). To a lesser extent, a similar trend was seen with FK506 alone or in combination with bupropion. Interestingly, the two drugs currently used in the clinic to treat obesity, bupropion and naltrexone, did not induce weight loss, either alone or in combination, in the obese rats fed a high fat diet. Overall, our data show temporary anorexigenic efficacy of sibutramine and in tendency FK506 ± bupropion.

### Food intake, but not the composition of the diet, is influenced by treatment with weight-loss associated drugs

Based on our finding of weight loss by the known oral anorexiant and appetite suppressant sibutramine, we next asked whether the rats showed changes in their preference and overall consumption of food in response to the various drug treatments. Rats were granted ad libitum access to three different pots containing chocolate, peanuts or powdered High Fat Diet (HFD). All groups showed a comparable preference for chocolate, making up to 40-50% of their daily intake, followed by HFD and peanuts (Figure 2A).

**Figure 2.**
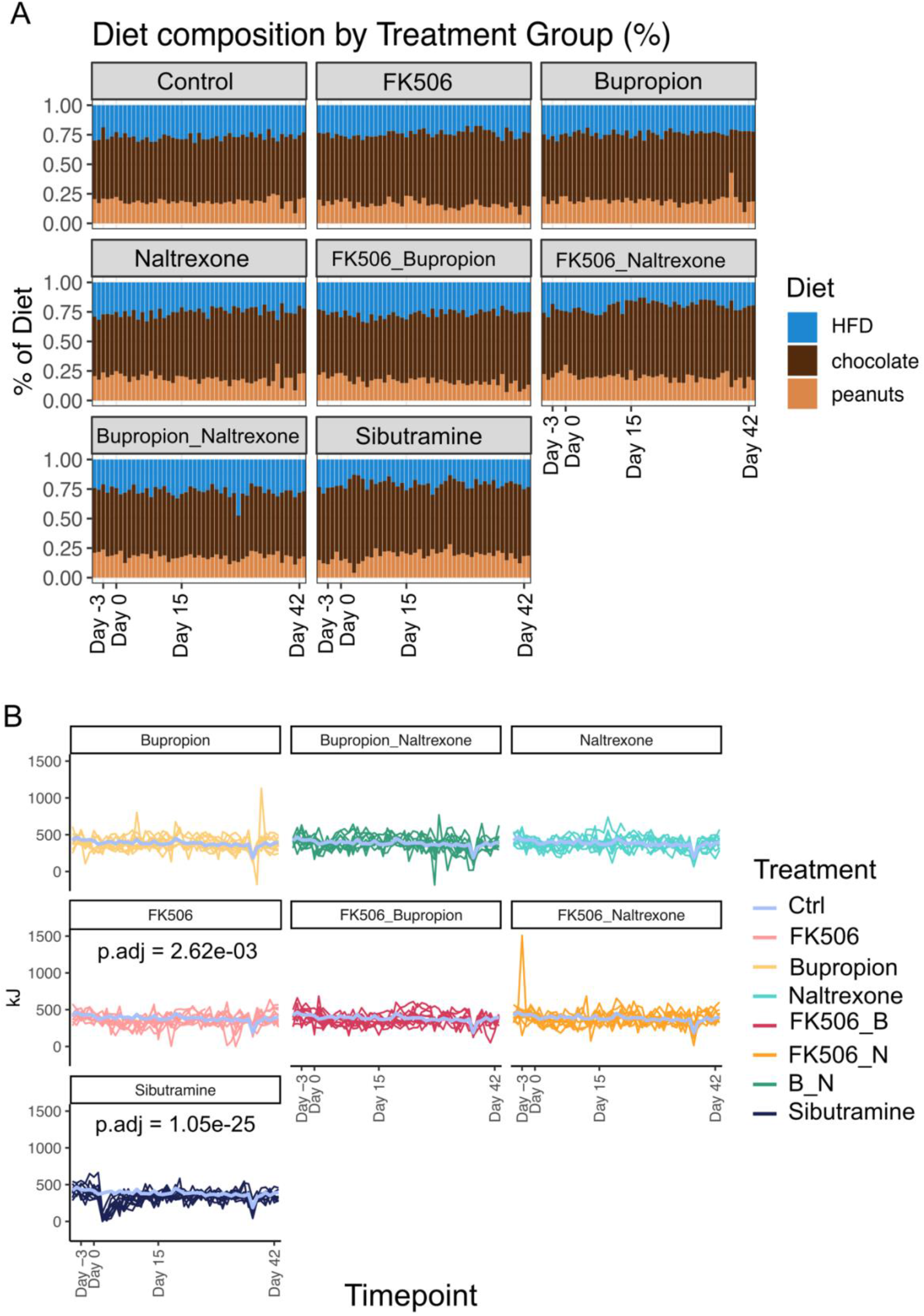
Food intake in obese rats treated with anti-obesity drugs. A) Bar plot representing the percentage of high fat diet (HFD), chocolate or peanuts consumed by each rat throughout the experiment. B) Combined food intake (kJ) for each rat throughout the duration of the treatment. A violet line represents the average food intake for rats in the control group. Statistical significance was assessed by ANOVA with the model: food intake value ~ Day + Treatment Group + Day*Treatment Group + sample, (function anova_test within the rstatix R package, parameters: wid = sample, within = Day, between = treatment, dv = food intake value) [74], and significant adjusted p-values for the day by group statistic have been reported on the appropriate panels.

FK506 treatment led to a moderate but significant decrease in food consumption (P.adjusted = 2.62e-03, Figure 2B, Table 3). The decrease in overall food intake compared to vehicle was more profound for sibutramine, and mainly driven by a temporary decrease within the first week of treatment (P.adjusted = 1.05e-25, Figure 2B, Table 3). This temporary effect within the first few days of treatment, based on a statistical assessment by ANOVA with the model: value ~ Day + Treatment + Day*Treatment + sample, is best observed when significant adjusted p-values for the day by treatment statistic are compared on the appropriate panels in Suppl. Figure S2. Here, statistical significance for all treatment groups was also observed at Day 36, where rats were fasted in preparation for the oral glucose tolerance test (OGTT) (see Supplementary Figure S2). Furthermore, both sibutramine and FK506 have a significant association of Day by Treatment, but inspection of their generalized effect sizes (GES) revealed that sibutramine treatment explained 19% of the variance observed, while FK506 treatment 7%. Overall, the effects of sibutramine and FK506 on weight loss in obese rats can, at least in part, be explained by a reduction in food consumption without any changes in food preference.

### Sibutramine, and FK506 in combination with bupropion, improve insulin secretion and reduce blood glucose levels in obese rats

Next, we explored whether weight loss or other aspects of the drug treatments would influence the response to glucose. Insulin resistance is common in obese patients, which can lead to the development of type 2 diabetes [8] and high blood glucose levels. Rats were subjected to an oral glucose tolerance test (oGTT) towards the end of the treatment (Day 36) and blood samples were taken before (−60, 0) and after (10, 20, 30, 45, 60 and 120 minutes) the oral challenge with 2 g glucose per kg body weight. Decreased excursions of plasma glucose suggested improved glucose tolerance in response to FK506 alone, in combination with bupropion or with naltrexone, compared to vehicle controls (Figure 3A). We moreover observed decreased insulin excursions for sibutramine, FK506, FK506 with bupropion and FK506 with naltrexone, compared to vehicle controls. This concomitant decrease in blood glucose and insulin levels during the oGTT points to an improved glucose tolerance and insulin sensitivity in rats treated with sibutramine. The impact of FK506 or FK506 with naltrexone on insulin excursions, or FK506 with naltrexone or bupropion on glucose excursions, was less clear, and could be based on beneficial effects on glucose tolerance and insulin sensitivity, but also a detrimental impairment of insulin secretion.

**Figure 3.**
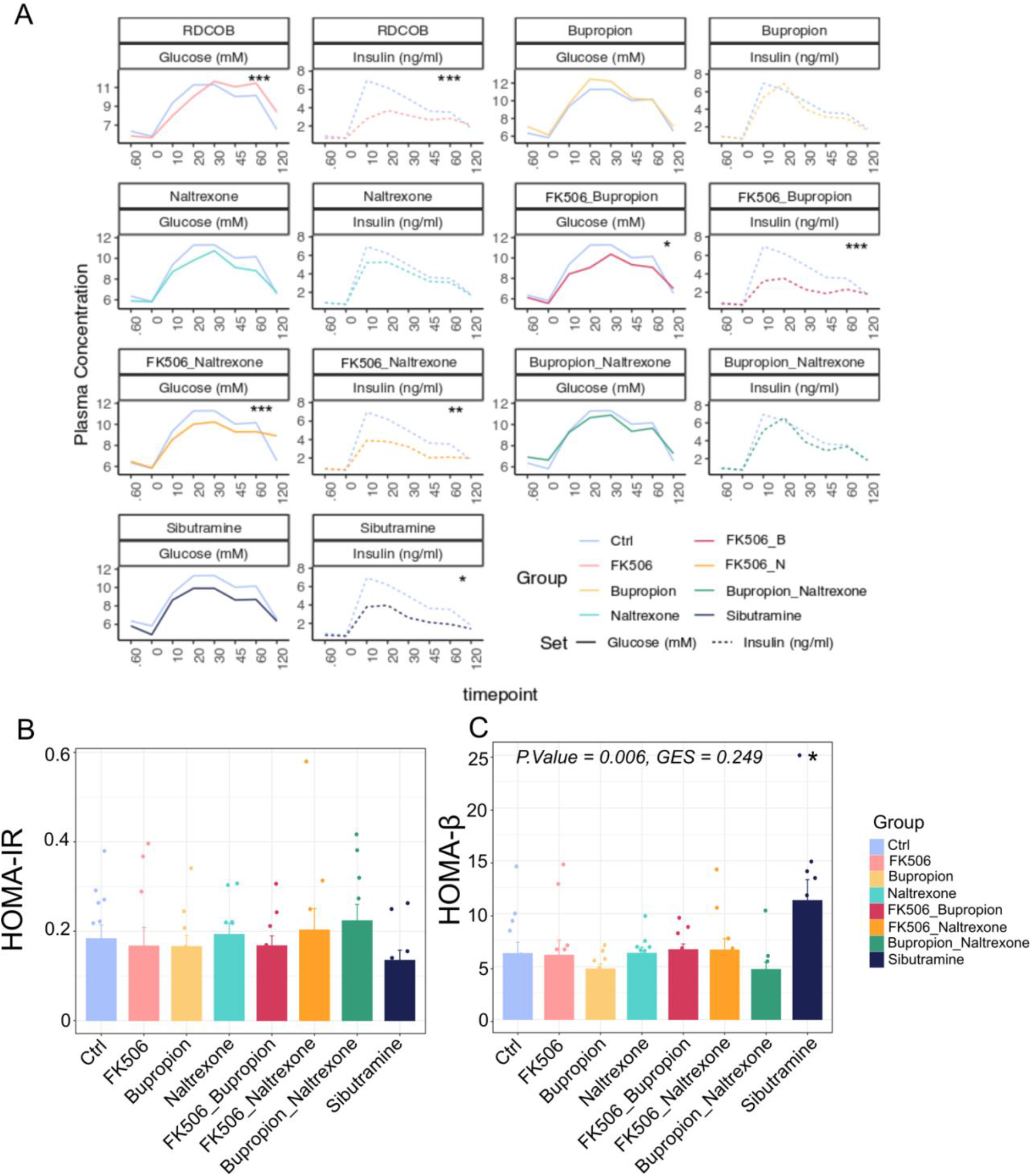
Assessment of glucose homeostasis in obese rats treated with anti-obesity drugs. A) Line plot of the treatment group average values for plasmatic Glucose and Insulin throughout the oGTT. Each treatment is plotted against a violet line, representing Glucose and Insulin levels in the controls. Statistical significance was assessed by ANOVA with the formula: value ~ Treatment Group + timepoint + timepoint * Treatment Group + sample, (function anova_test within the rstatix R package, parameters: wid = sample, within = Day, between = treatment, dv = value) [74]. P.values were corrected using the Benjamini-Hochberg method (Adjusted p.values: FK506, Glucose = 9.69e-04, Insulin = 1.98e-04; FK506_Buproprion, Glucose = 0.022, Insulin =4.17e-04; FK506_naltrexone, Glucose = 4.46e-04, Insulin = 7.41e-03; sibutramine, Insulin = 0.0256). Statistically significant results have been printed on the corresponding panels. Statistical significance levels = ns P > 0.05; * P < 0.05; ** P < 0.01; *** P < 0.001 (Control: N=12; Treatment Groups: N=10 rats per group). B) Average HOMA-IR values for each treatment group. Fasting insulin and blood glucose measurements were taken from the 0 timepoint of the oGTT test, then HOMA-IR values were calculated according to the formula: (fasting insulin [ng/mL] × fasting blood glucose [mg/dL])/405 [46]. Data represented as mean ± SEM. Statistical significance was assessed by one-way ANOVA test using the model: HOMA-IR ~ Group + Cohort (P.value = 0.685, GES = 0.062, n.s). C) Bar plot representing the values of HOMA-β, as function of insulin secretion and β cells activity, calculated with the formula: HOMA-β =(20× fasting insulin [μU/ml])/(fasting glucose [mmol/L] −3.5)[47]. Data was taken from the 0 timepoint in the oGTT and is represented as mean ± SEM. Statistical significance was assessed by one-way ANOVA test using the model: HOMA-β ~ Group + Cohort (Adjusted P.value = 0.006, GES = 0.249, **), followed by Tukey’s post-hoc test, which indicated the Ctrl vs. sibutramine comparison as statistically significant (Adjusted P.value = 0.0243, *). Statistical significance levels = nsP > 0.05; * P < 0.05; ** P < 0.01; *** P < 0.001 (Control: N=12; Treatment Groups: N=10 rats per group).

To further delineate the impact of our drugs on glucose control, we next applied the plasma glucose and insulin values of time point 0 from the oGTT to a homeostasis model assessment (HOMA) of insulin resistance (HOMA-IR) and beta cell function (HOMA-β) [46][47], and further assessed post-mortem pancreatic weights and insulin content in rats at Day 42. While HOMA-IR index values were comparable for the control rats and the treatment groups (Figure 3B), HOMA-β values were significantly higher in sibutramine-treated rats, indicating improved β cell function and insulin secretion (Figure 3C). In contrast, unchanged pancreatic weights but a significant decrease in insulin content in all rats treated with FK506, either alone or in combination (Supplementary Figure S3), suggest detrimental effects on the pancreatic production of insulin, an effect severely limiting the therapeutic potential of FK506 as weight loss agent [46,48].

Of the drugs tested, only sibutramine and FK506 in combination with bupropion lead to an overall reduction in weight in the cohort and improved insulin sensitivity in the oGTT assay. These physiological changes were associated with reduced food intake in response to sibutramine treatment, but not for the treatment with FK506 in combination with bupropion, suggesting distinct mechanisms. FK506 treatment was, moreover, associated with decreased pancreatic insulin content, an effect likely pointing toward unwanted side effects and limited potential as weight loss treatment.

### Impact of weight loss drugs on the microbiome

To assess whether the weight loss drugs affected the microbiome, we initially performed diversity analyses on the sequences obtained. α diversity measures the diversity of microbial species within an individual and it is generally lower in obese patients compared to their lean counterparts [49]. We found no statistically significant changes in alpha diversity at Day 42 vs. Day −3 for any of the treatments investigated (Figure 4A).

**Figure 4.**
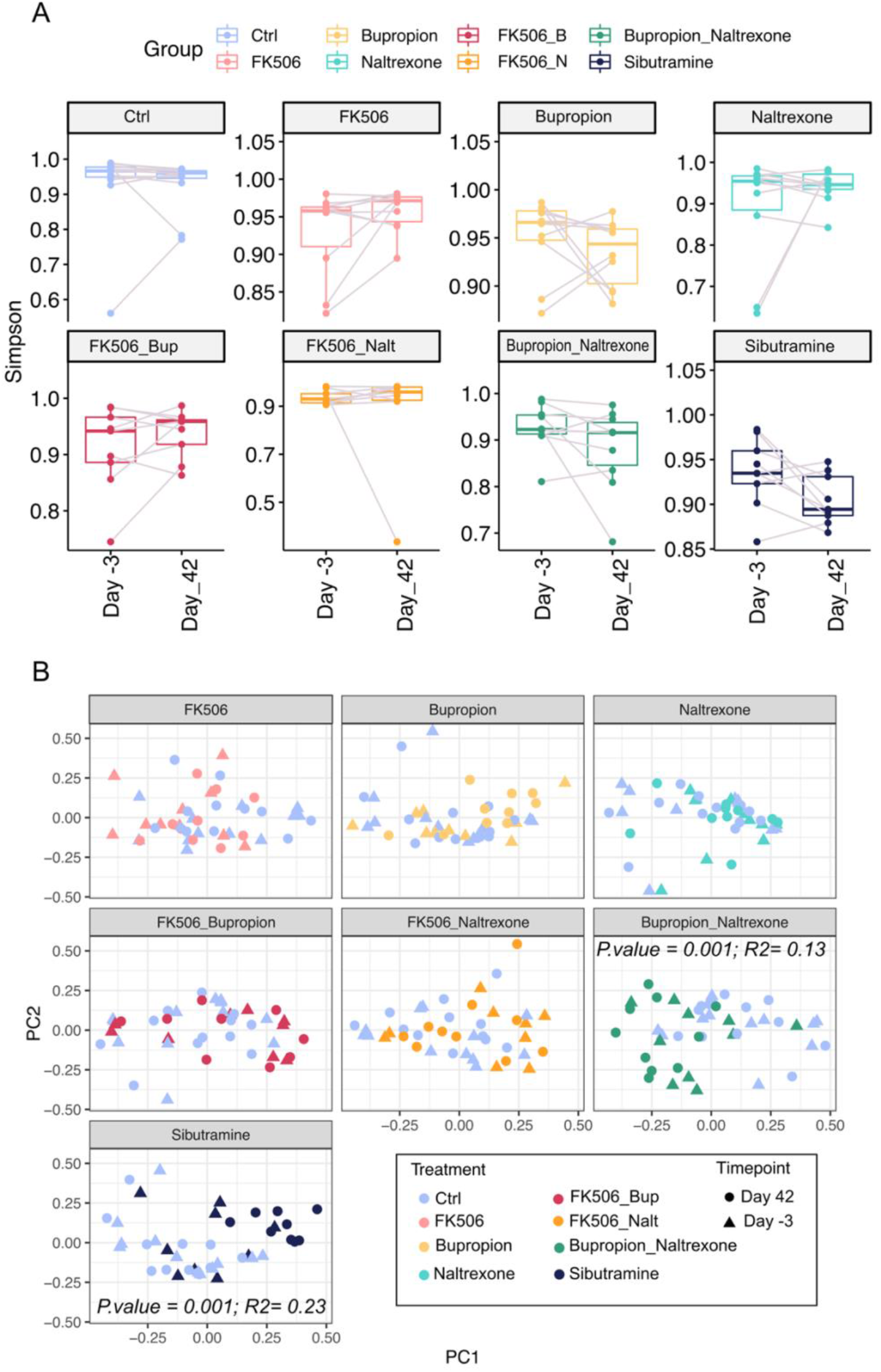
Diversity analyses on all treatment groups reveal no clear alteration of alpha diversity but some differences in beta diversity. A) Boxplots of Simpson’s index to estimate a diversity changes in each treatment group between the start (Day -3) and the end of the treatment (Day 42). Diversity analysis was carried out using Phyloseq’s estimate richness function. Statistical significance was calculated by Student’s paired t-test and resulting P.values were further corrected by the Benjamini-Hochberg method, but none of the comparisons were statistically significant. Notably, the higher the alpha diversity, the lower the Simpson index value will be. B) PCA plots showing β diversity amongst samples, calculated on relative abundances by Bray-Curtis distance between each sample, using the phyloseq R/Bioconductor package [38]. Statistical significance was assessed by the adonis test (Treatment * Group + cohort). The combination of bupropion and naltrexone, as well as sibutramine, were statistically significant. bupropion and naltrexone: P.value = 0.001, R^2^= 0.13, sibutramine: P.value = 0.001, R^2^= 0.23.

β diversity assesses the diversity in species composition between samples. Species composition was altered by sibutramine (R^2^ = 0.23, p.value = 0.001), or a combination of bupropion and naltrexone (R^2^ = 0.13, p.value = 0.001), but no other drug or combination (Figure 4B). A principal component analysis (PCA) of β diversity from the different cohorts of sibutramine-treated rats showed control samples clustering with samples taken before the start of the treatment (Pre), while samples from sibutramine treated rats fell into a separate cluster (Figure 5A). Furthermore, ordination analysis of the taxa present in our dataset highlighted a group of Bacteroidetes species separating along the first principal component (PC1), while a smaller cluster of Clostridiales spread across PC2 (Figure 5B).

**Figure 5.**
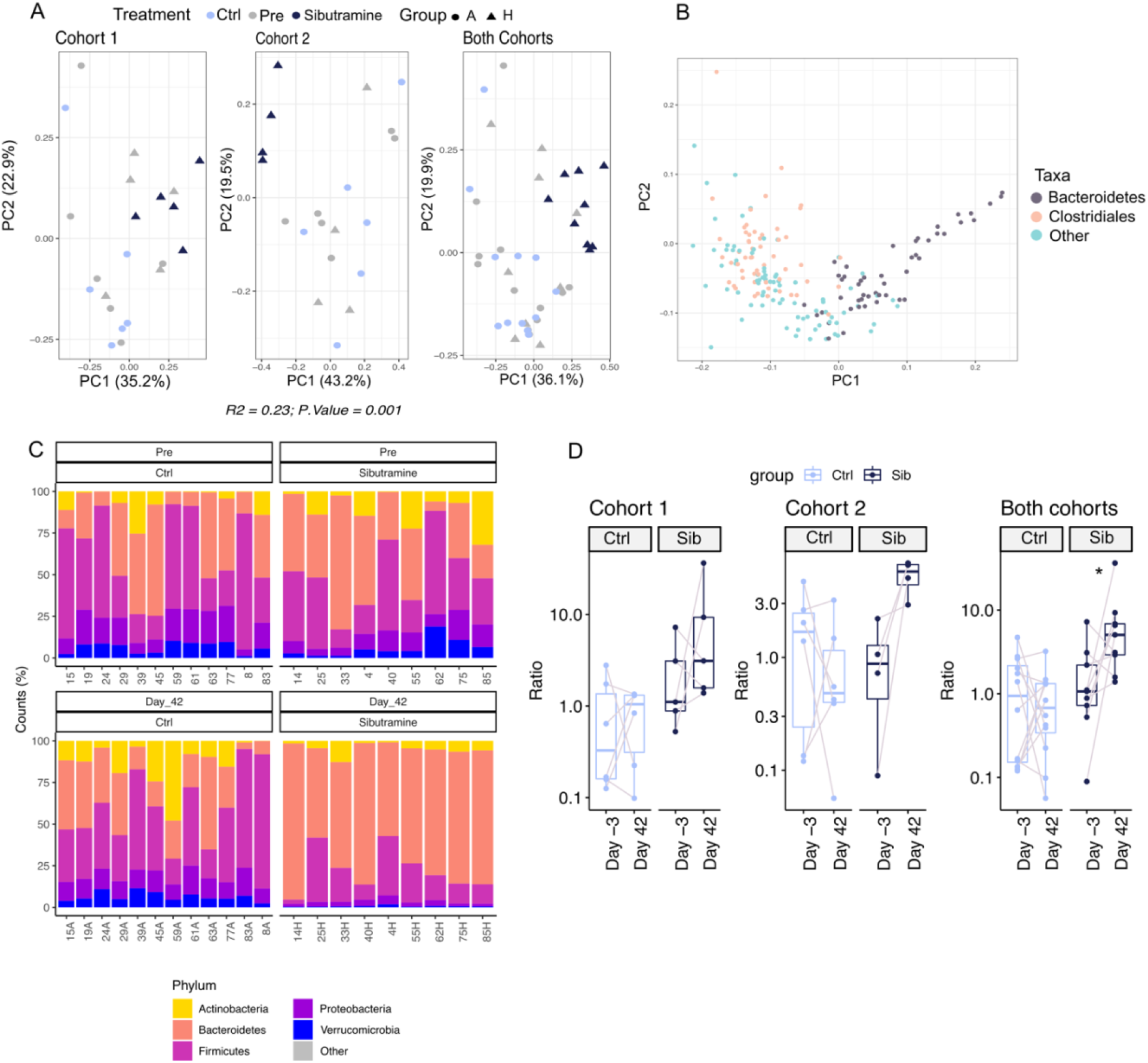
Sibutramine treatment impact on the microbiome. A) PCA of β diversity amongst samples, calculated on relative abundances by Bray-Curtis distance between each sample, using the phyloseq R/Bioconductor package [38]. Statistical significance was assessed by adonis test (Treatment * Group + cohort) and resulting variance explained (R^2^) and p.value are printed at the bottom of the plot. Plot shapes describe the different treatment groups, with circles representing the Control group (A) and triangles the sibutramine-treated samples (H). B) PCA of β diversity amongst taxa, calculated on relative abundances by Bray-Curtis distance between each sample, using the phyloseq R/Bioconductor package [38]. C) Bar plot summarizing the percent composition of the top 5 phyla across control (left panel) or sibutramine (right panel) samples, at the start (top) or the end (bottom) of the treatment. D) Boxplots of the comparison of Ratio Bacteroidetes/Firmicutes between Day −3 and Day 42 of Ctrl and sibutramine-treated group. Statistical significance was assessed by T-test. Statistical significance levels: ns P > 0.05; * P < 0.05; ** P < 0.01; *** P < 0.00 (Control: N=12; sibutramine: N=10 rats per group).

Using the currently available annotations of the bacterial species and their genes, we were unable to gain any mechanistic insight into how the combination treatment of bupropion and naltrexone might be affecting the host (Supplementary Figures S4 and S5). However, sibutramine treatment changed the microbiome in ways consistent with weight loss in the obese rats and was selected for more detailed analysis.

### Sibutramine treatment increases the Bacteroidetes to Firmicutes ratio

We next evaluated whether treatment with sibutramine caused a shift in the ratio between Firmicutes and Bacteroidetes (Figure 5C and 5D). Several lines of evidence suggest that in obese individuals, microbes belonging to Firmicutes phylum are generally enriched compared to Bacteroidetes, thus giving a lower ratio of Bacteroidetes to Firmicutes [10,18,50]. Upon weight loss, in both cohorts sibutramine treated rats had an increased Bacteroidetes to Firmicutes ratio at day 42 vs. day −3 (sibutramine mean at Day −3: cohort 1 = 3.04, cohort 2 = 1.07, both cohorts = 2.16; sibutramine mean at Day 42: cohort 1 = 12.23, cohort 2 = 5.75, both cohorts = 9.35).

### 192 species change on sibutramine treatment

We next focused on the species differences between the gut microbiome of sibutramine-treated rats before (Day–3) and after sibutramine treatment (Day 42). Analysis of differentially abundant species between the two groups highlighted statistically significant changes in 192 species (Figure 6). Amongst these, thirteen different species belonging to the Bacteroides genus were significantly increased by Day 42; *B.thetaiotamicron, B.caecimuris, B.heparinolyticus, B. cellulosilyticus, B. fragilis, B.caccae, B.vulgatus, B.salanitronis, B.ovatus, B.helcogenes, B.zoogleoformans, B.dorei*. Bacteria of this genus, and more generally of the Bacteroidetes phylum, have been found decreased in obese individuals in favour of Firmicutes species [14,50]. At the phylum level, Bacteroidetes were significantly increased by Day 42, while Firmicutes did not show overall significant changes, perhaps due to the fact that some of their species were increasing, while others were decreasing. In addition, Actinobacteria species detected showed a significant decrease by the end of the treatment (Figure 6A, green panel).

**Figure 6.**
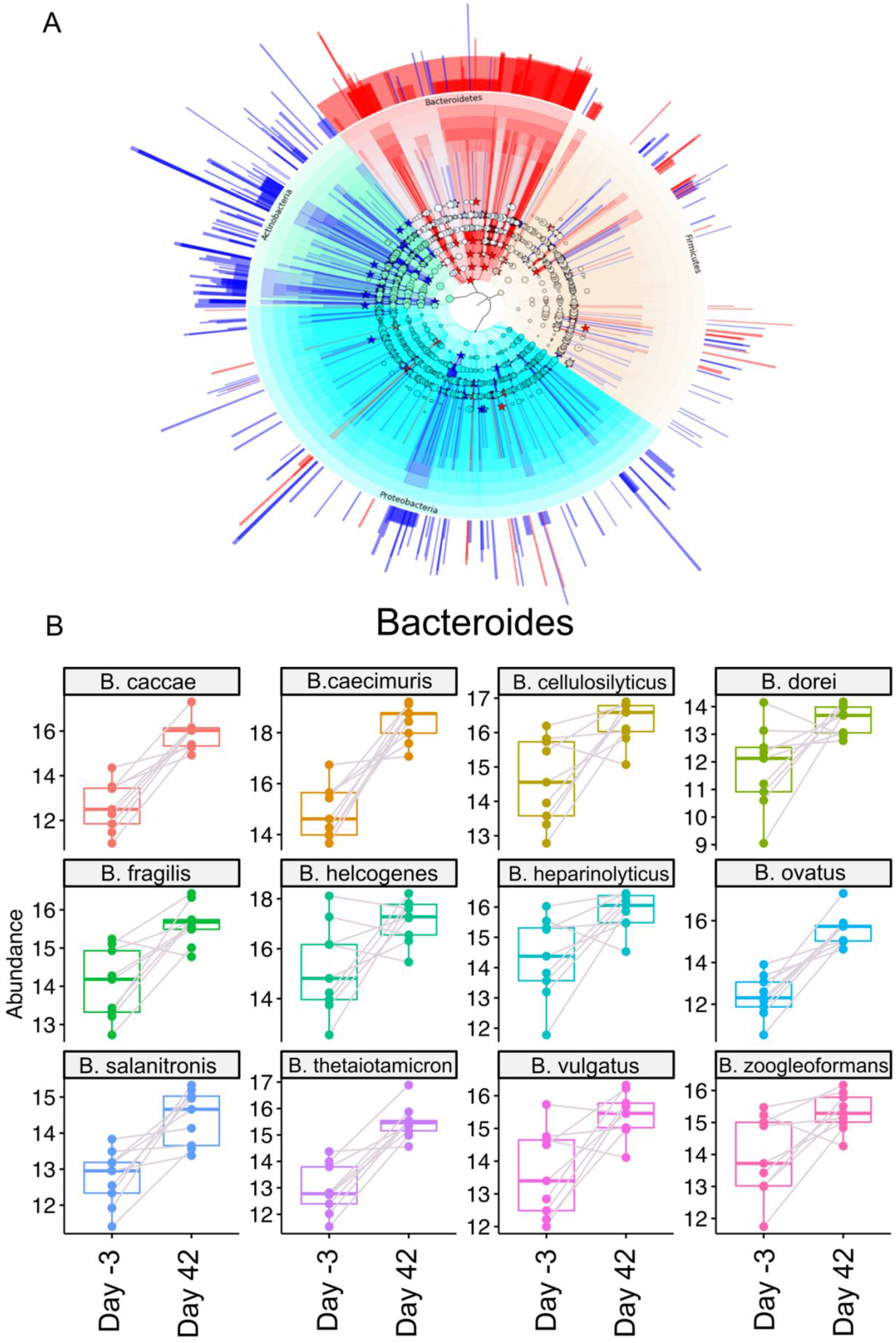
Differentially enriched species between rats before (Day -3) and after (Day 42) sibutramine treatment. A) Phylogenetic tree representation of differentially expressed taxa between samples at the start and end of the treatment. Differential enrichment at every classification level (Phylum, Class, Order, Family, Genus and Species) was calculated with R package DESeq2 using a linear model (~ Day + rat number). P values were adjusted by Benjamini-Hochberg correction and only species with P.adj <= 0.05 were considered significantly different, then tables with log2 and library size normalized counts (rlog) for every taxon were extracted [41,73]. These were fed into GraPhlAn for phylogenetic tree visualization. External annotation corresponds to Log2 fold-change values for every differentially expressed species. Species of interest were indicated in the circular tree by stars and coloured according to their Log2 Fold change (Log2 fold-change < 0, blue; Log2 fold-change > 0, red). Concentric circles each represent a taxonomical level, starting from Phylum (Proteobacteria, light blue; Firmicutes, yellow; Bacteroidetes, red; Actinobacteria, green), then proceeding outwards to Class, Order, Family, Genus and Species. Notably, Bacteroidetes as a whole are significantly increased by the end of the treatment (see red star in the innermost circle). B) Boxplots showing levels of differentially enriched Bacteroides species, before and after sibutramine treatment. Grey lines connect dots corresponding to the same rat. Counts used were log2 transformed and normalized by library size [73].

Moreover, a redistribution of several probiotic species is evident. Five members of Bifidobacteriaceae, such as *B.pseudolongum, B.animalis, B.catenulatum, B.dentium* and *B.scardovii* were significantly decreased, while two Lactobacillales, *L.reuteri and L ruminis*, increased by the end of the treatment. Taken together, results show that sibutramine treatment significantly alters microbiome composition in obese rats.

### Functional analysis of microbiomes reveals significant decrease of “Flagellar Assembly” related genes

Single species changes might not explain all the differences in microbiome composition before or after sibutramine treatment. An advantage of shotgun metagenomics is the possibility of sequencing the gene pool of the microbiome, rather than just the 16S rRNA, thus allowing the detection of individual genes with the potential to be expressed. Analysis of differentially abundant genes revealed statistically significant changes in 1047 genes, of which 536 are increased and 509 decreased in sibutramine-treated rats at the end of the treatment compared to the beginning (Figure 7A). To functionally characterize these genes, we performed gene set enrichment analysis (GSEA). This method aims at finding enrichment of entire gene sets rather than single genes, thus providing insight into the biological pathways contributing to a particular phenotype [51]. Results show a significant enrichment of genes from the “Flagellar Assembly” set at the lower end of the ranked list, indicating their decrease in sibutramine-treated samples (Normalized enrichment score (NES) = -1.71, p.adjusted = 0.03, Figure 7B). In bacteria, the flagellum is a hair-like structure used for locomotion (Figure 7C), and has been shown to induce inflammation due to stimulation of interleukin 8 (IL8) release from intestinal epithelial cells [52]. Moreover, different studies have shown an enrichment of this pathway in obese patients, as well as in patients with metabolic syndrome [50,53]. Further analysis of genes associated with this pathway highlighted a panel of 20 genes, all of which are differentially abundant and significantly decreased by the end of the treatment (Figure 7D).

**Figure 7.**
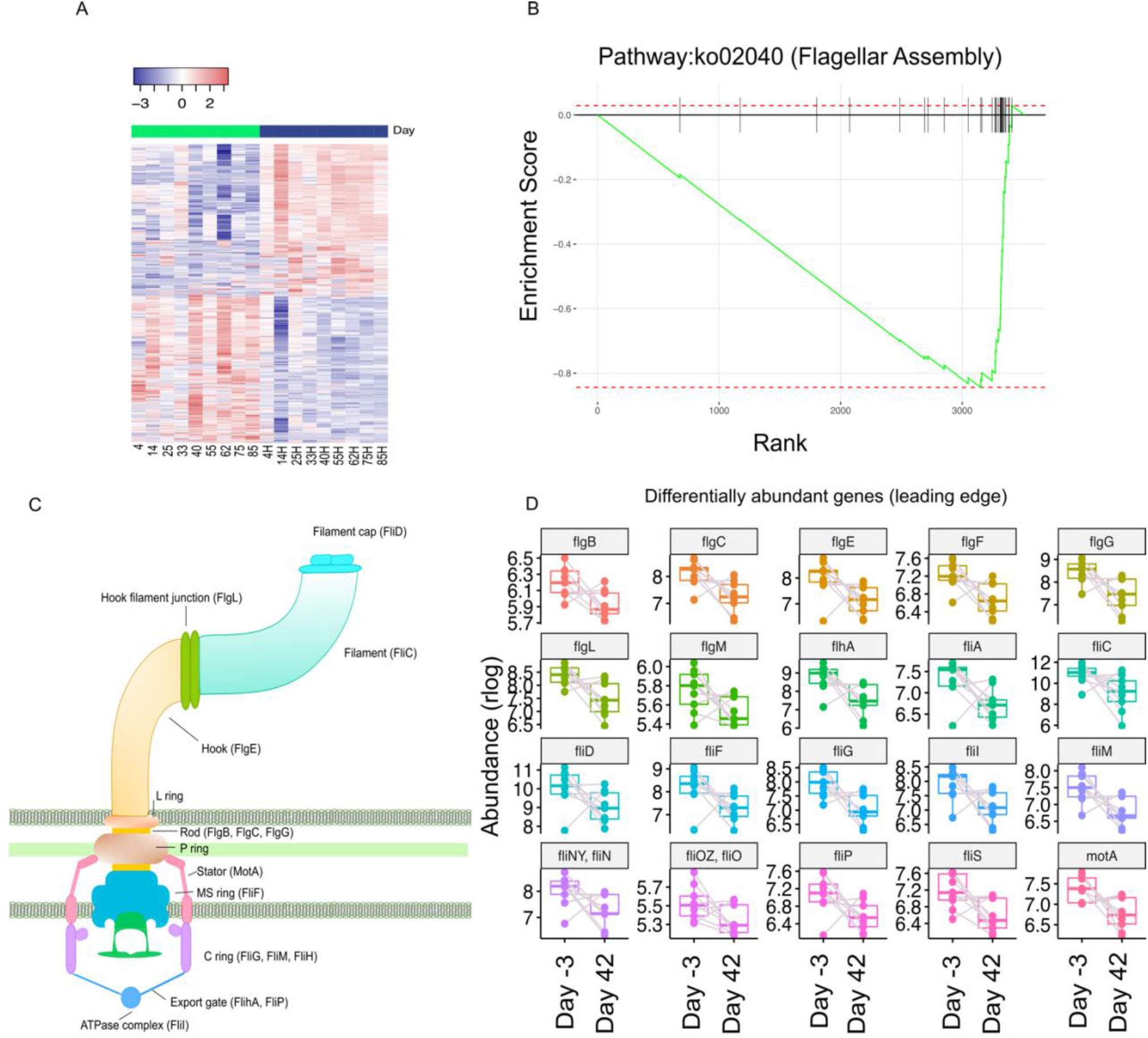
Gene Set Enrichment Analysis highlights decrease of genes related to flagellar assembly. A) Differential gene abundance between sibutramine-treated rats at the start and end of the treatment. Heatmap representing log2-transformed and normalized read counts for each of the 1047 differentially abundant genes. As with species, significant changes in gene levels were calculated using DESeq2 with the model ~ rat number + Day [73]. P values were corrected with the Benjamini-Hochberg method, and only genes with adjusted P value <= 0. 05 were considered statistically significant. Top bar shows which columns correspond to Day -3 (green) or Day 42 (blue) samples. B) Enrichment plot for the pathway “Flagellar Assembly” (ko:02040). Briefly, genes have been ranked by log2 fold change multiplied by the −logl0(p.adjusted), then gsea was performed with the fgsea R/Bioconductor package [72]. “Flagellar Assembly” was the only statistically significant result (normalized Enrichment Score: −1.7, p.adjusted = 0.03). C) Schematic representation of the bacterial flagellum and its protein components. Genes detected in our analysis are indicated on the figure. [54]. D) Boxplots of leading edge genes belonging to this pathway, all of which are differentially abundant in sibutramine samples at Day 42 compared to Day −3. Boxplots were obtained by plotting the log2-normalized expression values for each gene.

Genes in this panel encode for all the different components of the flagellum. *FliD, fliC, flgL* and *flgE* gene products make up for the extracellular portion of the flagellum, encoding for the filament cap, the filament itself, the hook junction and the hook, respectively [54]. *FlgB, flgC, flgF* and *flgG* encode for the rod, while *motA* and *fliF* are representative of the membrane-spanning basal body, encoding for a part of the rotor and the MS ring, which provides the base for flagellar assembly [55,56]. Finally, five genes, *fliM*, *fliN*, *flhA* and *fliP*, encode for proteins making up the export apparatus on the cytosolic side [57] (Figure 7C).

Further analysis of the species that are likely to contribute to the enrichment of three representative genes, one for each major flagellum component, *flgE*, *fliC* and *motA* (Figure 8A), found *Flavonifractor plautii* and *Roseburia hominis* as the two major sources of the respective genes (Figure 8B-D). Interestingly, while *F.plautii* is differentially enriched at Day 42, *R.hominis* levels do not significantly change over time *(F.plautii:* Log2-fold change = 1.86, p.adjusted = 0.03; *R. hominis:* Log2 fold change = 0.93, p.adjusted = 0.49).

**Figure 8.**
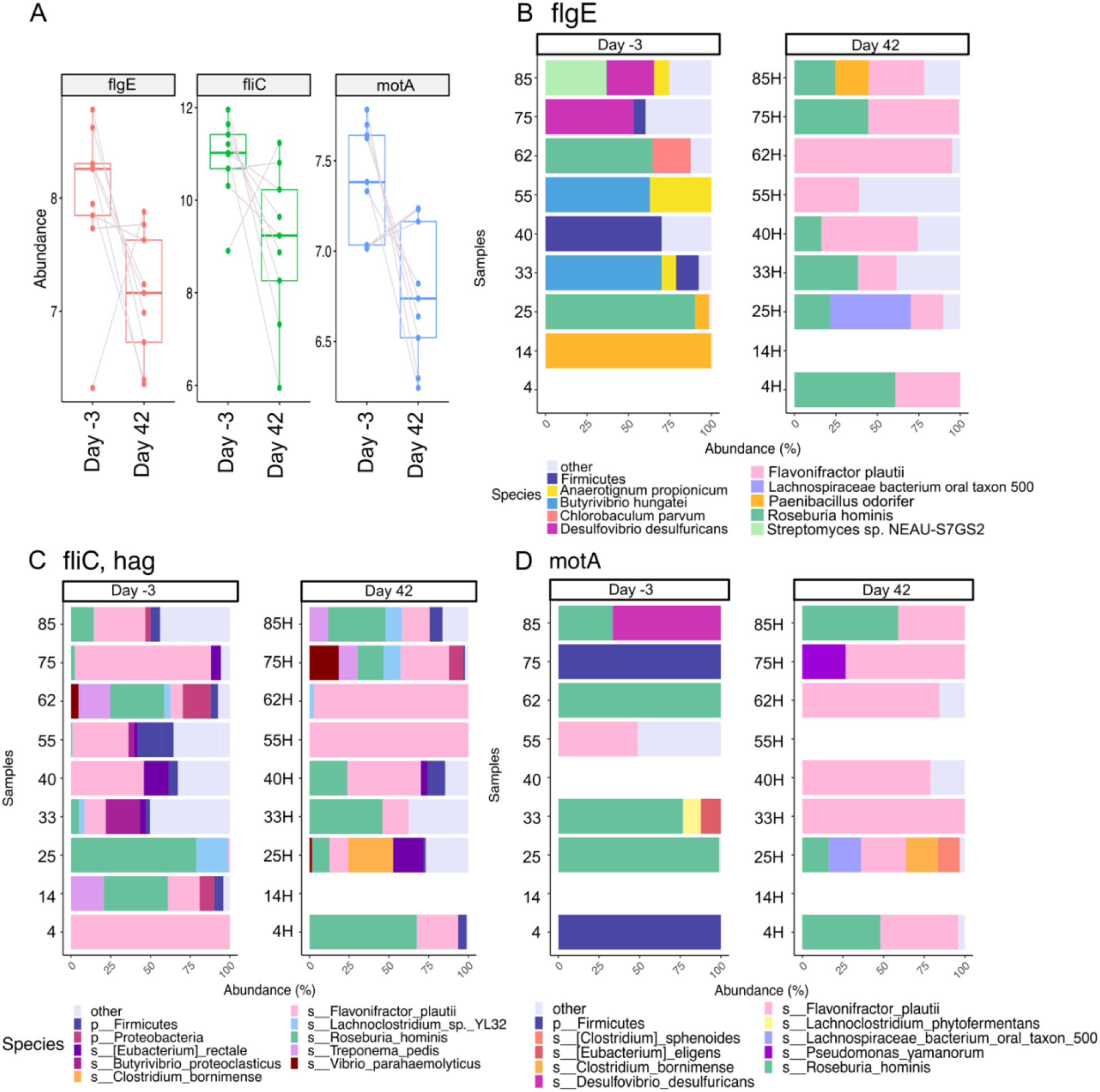
Species composition of three representative genes from the “Flagellar Assembly” pathway. A) Boxplots showing expression levels of three representative members of the “Flagellar Assembly” pathway. Expression values have been log2 normalized as in Figure 7. B-D) Species contribution to the levels of flgE (B)fliC (C) and motA (D). Gene counts for each species are expressed as percentage and only the top 10 species for each gene are represented, whilst the remainder have been collapsed under the “other” category. Empty rows correspond to samples that either had no reads annotated to the gene (motA, 55H and 14H; flgE 4 and 14H) or whose reads were entirely annotated to “k_unassigned” (fliC, 14H; motA, 14 and 40). Notably, every sample had a certain proportion of reads annotated to “k_unassigned” (Kingdom unassigned), and thus lacked a reliable annotation. These reads have been excluded from the dataset prior to plotting of fliC, flgE and motA. The percentage of unassigned reads for each gene, within every sample, is described in Table S2.

*R.hominis* is a butyrate-producing bacterium from the Firmicutes phylum, and while its impact on *flgE* counts seems to be increasing by Day 42, in both *motA* and *fliC* it remains constant throughout the treatment (Figure 8B). *F.plautii* is a flavonoid-metabolizing species from the Clostridiales family (Firmicutes). Its contribution to gene counts of *flgE* and *motA* increases in sibutramine-treated rats by Day 42 (Figure 8B and D), while it is always present in the counts of *fliC* (Figure 8C). Interestingly, this species has been described as enriched in lean individuals compared to obese patients and has been linked to anti-inflammatory activities, potentially suggesting a positive change in the rats microbiome induced by sibutramine treatment [58,59].

Thus, results from our functional analysis support a role of sibutramine treatment in positively affecting the gut microbiome. A decrease in genes related to flagellar assembly, previously associated with obesity-induced inflammation, as well as a higher contribution of anti-inflammatory species, such as *F.plautii*, to their differentially abundant gene counts suggests a shift towards a decreased inflammatory state, an opposite trend compared to patterns commonly associated with obesity.

## Discussion

Multiple reports suggested a functional impact of gut microbiome perturbations on the development of obesity and comorbid sequelae [18,60]. Similarly, numerous studies revealed that weight loss by dietary or surgical interventions was able to reverse perturbations in gut microbiome [61–63]. However, few studies to date analysed the specific impact of pharmacological weight loss agents on gut microbiome diversity. In our detailed analysis based on shotgun metagenomics, we reveal that the treatment of female rats with the appetite suppressant sibutramine leads to concomitant weight loss, improved glucose tolerance and insulin sensitivity, a trend towards increased α diversity, a distinct β diversity pattern and a shift in the Bacteroidetes to Firmicutes ratio towards a higher proportion of Bacteroidetes.

Sibutramine is a known inhibitor of serotonin reuptake, and more than 90% of serotonin production occurs in the gut, regulated by several microbial species [64–66]. Our analysis shows an influence of sibutramine treatment on the microbiome of obese rats, but it cannot define whether this change is causing weight loss, or whether weight loss induced by sibutramine is causing the changes in the gut microbiome. Faecal microbiota transplantation (FMT) from sibutramine-treated rats to obese rats could answer this open question and reveal the functional role of such sibutramine-induced gut microbiome changes on obesity. Such studies should further entail the study of gut serotonin production as a potential mediator of this effect.

The most striking alteration in the gut microbiome in response to sibutramine treatment were linked to the decrease of genes related to flagellar assembly. Flagella are rod-like structures composed by flagellin and used by bacteria for locomotion. As host toll like receptor 5 (TLR5) recognises flagellin and is able to activate an immune response, increased presence of motile bacteria in the gut microbiota has been liked to a pro-inflammatory state in a range of diseases, including metabolic syndrome [67,68]. Thus, a decrease of genes coding for this protein, along with other components of the flagellum might be indicative of a reduced inflammation in the gut of sibutramine treated rats. As suggested by the significant decrease of genes involved in flagellar assembly, differentially abundant genes hint at positive changes to the gut microbiome, although results must be interpreted carefully. Indeed, while recent research in the field has highlighted microbial patterns associated with obesity at higher classification levels, such as the ratio of Bacteroidetes to Firmicutes and a lower α diversity, many of the species significantly changing are still largely unknown. Another complication is that, while over time metagenomic databases have expanded, there is still a consistent portion of reads that remain unassigned. For these reasons, our results remain largely descriptive, mainly pointing toward the specific association of certain species, genes or pathways involved in flagellar assembly with weight loss induced by sibutramine treatment.

Between the beginning and the end of sibutramine treatment, we revealed a significant increase in 13 Bacteroides species, and in the Bacteroidetes phylum as a whole. In the context of obesity, Bacteroides species such as *B. thetaiotamicron*, were shown to be decreased in obese individuals where they contribute to glutamate fermentation and short chain fatty acid production [17]. Several studies linked an increased presence of Firmicutes species in obese individuals with improved efficiencies in extracting energy from nutrients and excessive storage [14,69]. Overall, our data in sibutramine-treated rats are thus consistent with the reported, beneficial shift from Firmicutes to Bacteroidetes, including *B. thetaiotamicron*. However, microbiome research remains a growing and controversial field, with many species remaining largely uncharacterised. Recent advances in sequencing techniques may help bypassing the limitation of culturing species in the lab and thus overcome our lack in knowledge on the functional impact of distinct species. Our complete shotgun metagenomics dataset will serve as resource and step in that direction, to ultimately support the study of functional roles of specific microbial species and genes in metabolic diseases such as obesity.

Unexpectedly, we observed considerable variability in the weight loss between and within the intervention groups. naltrexone and bupropion, contrary to earlier published reports, failed to induce significant weight loss, with naltrexone-treated rats gaining weight by Day 42 [24]. Reasons for this discrepancy remain elusive but may in part be based on our choice of using female Wistar rats and/or our dietary feeding regimen.

Our study also highlights other physiological changes to the obese rats that appear unrelated to weight loss or changes to the microbiome composition that might warrant further investigation, for example the impact of FK506 combined with bupropion or naltrexone on insulin sensitivity and blood glucose levels. In non-obese rats, FK506 is associated with maintenance of a healthy weight compared to untreated controls [21] (Supplementary Figure S1). In our obese rats, FK506 elicited at best modest effects on body weight and food intake but induced a profound decrease in insulin secretion upon a glucose challenge when given either alone or in combination with naltrexone or bupropion. Further investigation of pancreatic insulin levels suggested a detrimental effect of FK506 on insulin production in rats treated with this drug, alone or in combination. Overall, our data suggest that FK506 has a restricted therapeutic window, as obese patients often show insulin resistance and are at high risk of developing diabetes.

## Conclusions

Shotgun metagenomics analysis of the gut microbiome from obese female rats treated with a panel of weight-loss drugs revealed that common pharmacological treatments for obesity, such as bupropion and naltrexone, did not induce consistent weight loss, while sibutramine successfully reduced the weight of the rats by decreasing their appetite. Analysis of the rats’ gut microbiome highlighted a potentially beneficial impact of sibutramine treatment, as their microbiome composition by the end of the treatment is significantly different both compared to the beginning of the treatment and to control rats. Differential analysis revealed a significant increase in the abundance of species belonging to the Bacteroidetes phylum, and a concomitant shift of the Bacteroidetes to Firmicutes ratio towards the former, while in obese patients the ratio favours Firmicutes. Functional analysis showed a significant decrease in the abundance of genes coding for proteins of the flagellum, a bacterial component that has been linked to inflammation, further suggesting a beneficial impact of the treatment on the gut microbiome of obese rats.

These results lay the foundation for further investigation of the ways in which the transformed microbiome is affecting the host metabolism. To this end, a metabolomic analysis of the blood of sibutramine-treated rats would shed light on the kind of metabolites that the microbiome is producing, or sibutramine is inhibiting, leading to a better understanding of the connection between the gut microbiota and its host metabolism, in the context of obesity.

Furthermore, the entire shotgun metagenomic dataset is made publicly available as a resource for the field.

## Acknowledgements

This work was supported from the European Union’s Horizon 2020 research and innovation 260 Programme under the Marie Skłodowska-Curie Grant Agreement No. 675610 (Chromatin and Metabolism - ChroMe) and Chronos Therapeutics Ltd. Chronos Therapeutics Ltd. (Oxford, UK) funded the animal experiments conducted by RenaSci Ltd. (Nottingham, UK).

## Author Contributions

JS, KT, HJ and JM designed the animal experiments; SR sequenced and analysed the composition of the fecal extracts, and conducted the bioinformatic analysis with supervision from JM. NI, SR and JM interpreted the DNA sequencing data. PF, SR and JM interpreted the physiological data. SR and JM wrote the manuscript with input from other authors.

## Supplementary

**Figure S1.**
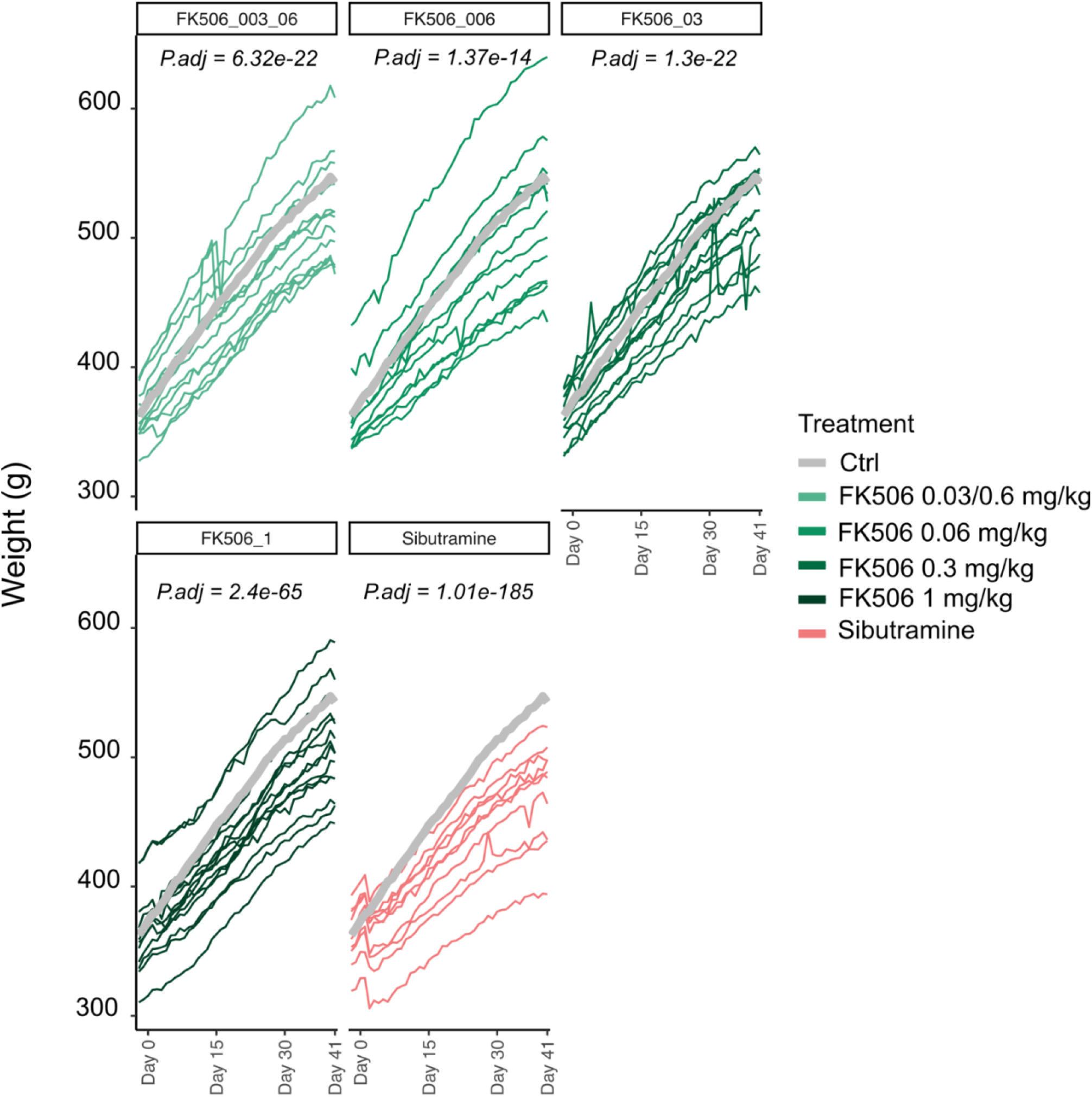
Long-term treatment of adult rats maintained on a high fat diet with different dosages of FK506 induces weight loss and promotes maintenance of a healthier weight. FK506 therapeutic potential was assessed by treating 75 adult, male Sprague-Dawley rats maintained on a high fat diet for 40 days at different concentrations of drug. Rats treated with FK506 showed decreased body weight compared to untreated controls. Statistical significance was assessed by ANOVA with the model: weight ~ Day + Treatment Group + Day* Treatment Group + sample, (function anova_test within the rstatix R package, parameters: wid = sample, within = Day, between = treatment, dv = weight) [74], and significant adjusted p-values for the day by treatment group statistic have been reported on the appropriate panels. Generalized effect sizes for the Day*Treatment Group comparison were as follows: FK506 0.03/0.6 mg/kg =0.18, FK506 0.06 mg/kg = 0.1, FK506 0.3 mg/kg = 0.18, FK506 1 mg/kg = 0.34, sibutramine = 0.70.

**Figure S2.**
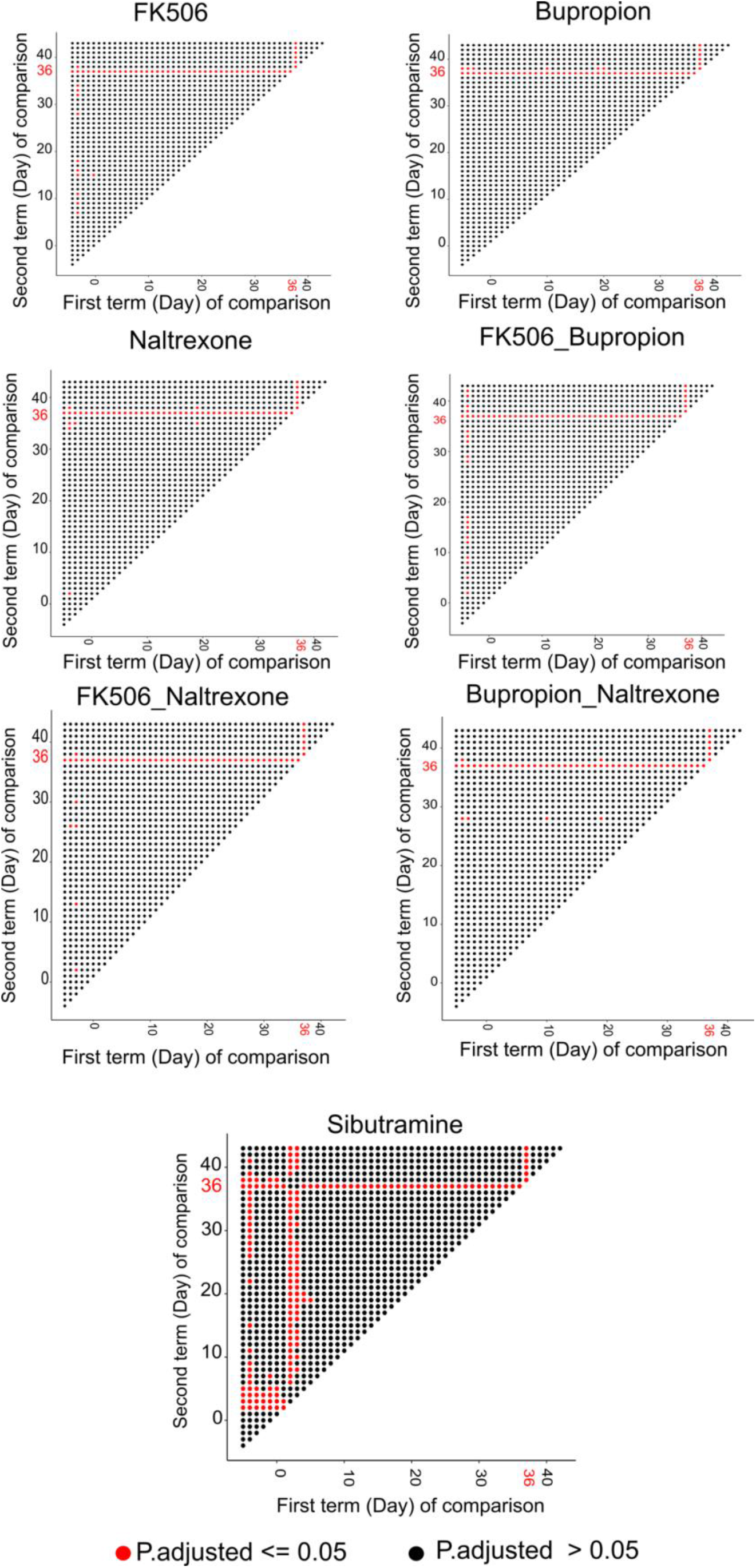
Tukey’s post-hoc test following ANOVA statistical test on timepoint reveals early but temporary anorexigenic effects of sibutramine, and significant changes due to fasting on Day 36. Visualization of the p.values (P.adj <= 0.05, red; P.adj > 0.05, black) in the Tukey’s post hoc test following ANOVA statistical test on timepoint. Food intake on a given day (first term of comparison, x axis) was compared to another (y axis) in the Tukey’s post-hoc test. The statistical test performed gives a significant p.value, indicating which days are contributing to the significance of the timepoint comparison. A closer look at the plots for most of the treatment groups, except sibutramine, reveals that timepoint is coming up as a significant term in ANOVA test due to the fasting occurred on Day 36, in preparation of the oGTT test.

**Figure S3.**
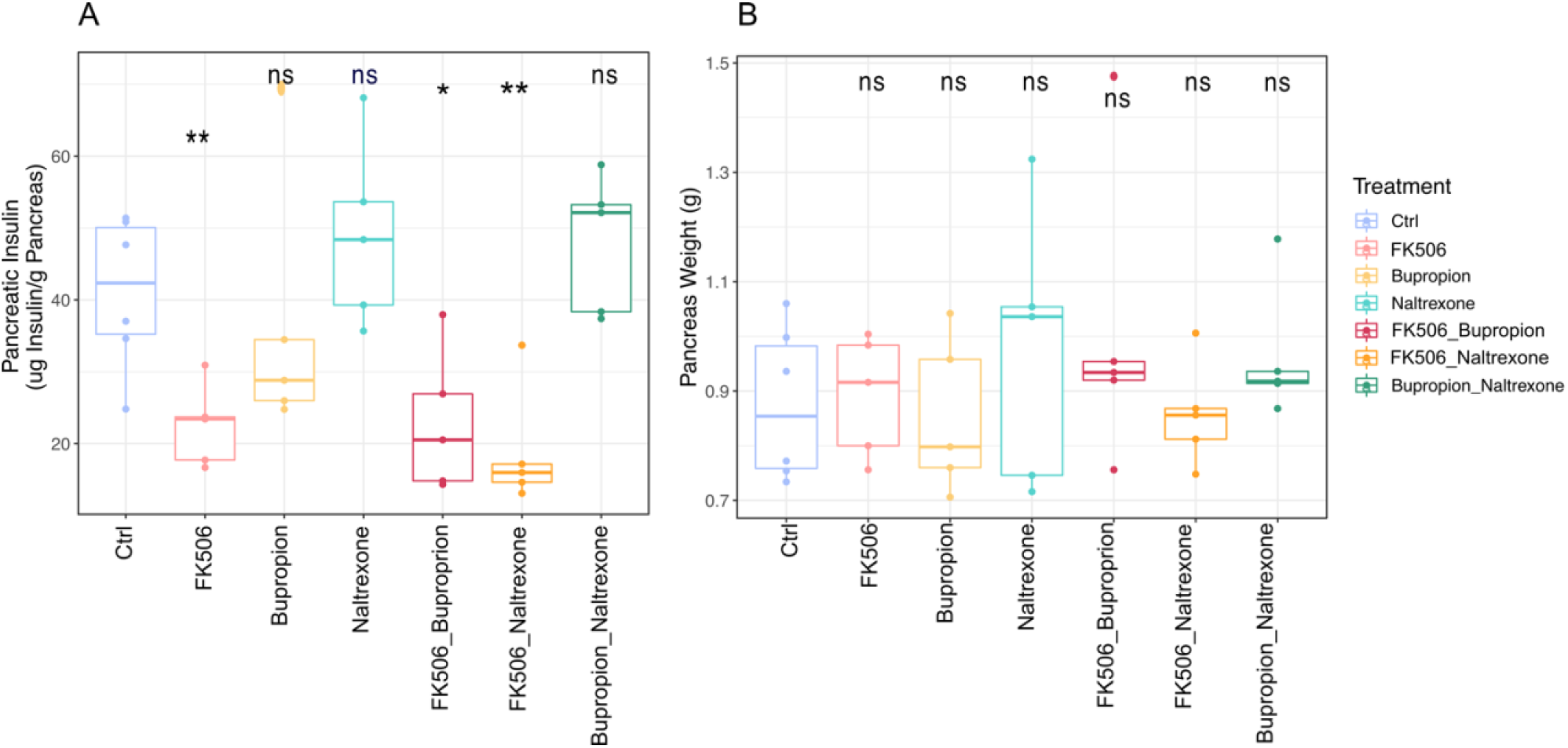
Pancreatic Insulin levels at Day 42. A) Boxplots representing pancreatic insulin levels at Day 42. Statistical significance was assessed both by Student’s T-test of each treatment group against Controls and by ANOVA. T-test detected a significant decrease in pancreatic insulin in all FK506-treated samples, either alone or in combination (P.adj: FK506 = 0.019, FK506_bupropion = 0.03, FK506_naltrexone = 0.019). Statistical significance was confirmed by ANOVA, calculated with the formula: pancreatic insulin ~ Treatment Group + Cohort (P.adj. = 0.001 (Treatment Group), 0.99 (Cohort)), but Tukey ‘spost-hoc test indicated no significance when comparing any treatment group against control. Significant comparisons have been described in Table S3. B) Boxplots of pancreas weight at Day 42. Statistical significance was assessed both by Student’s T-test and ANOVA (pancreas weight ~ Treatment Group + Cohort), but no statistical difference between controls and any of the treatment groups was detected.

**Figure S4.**
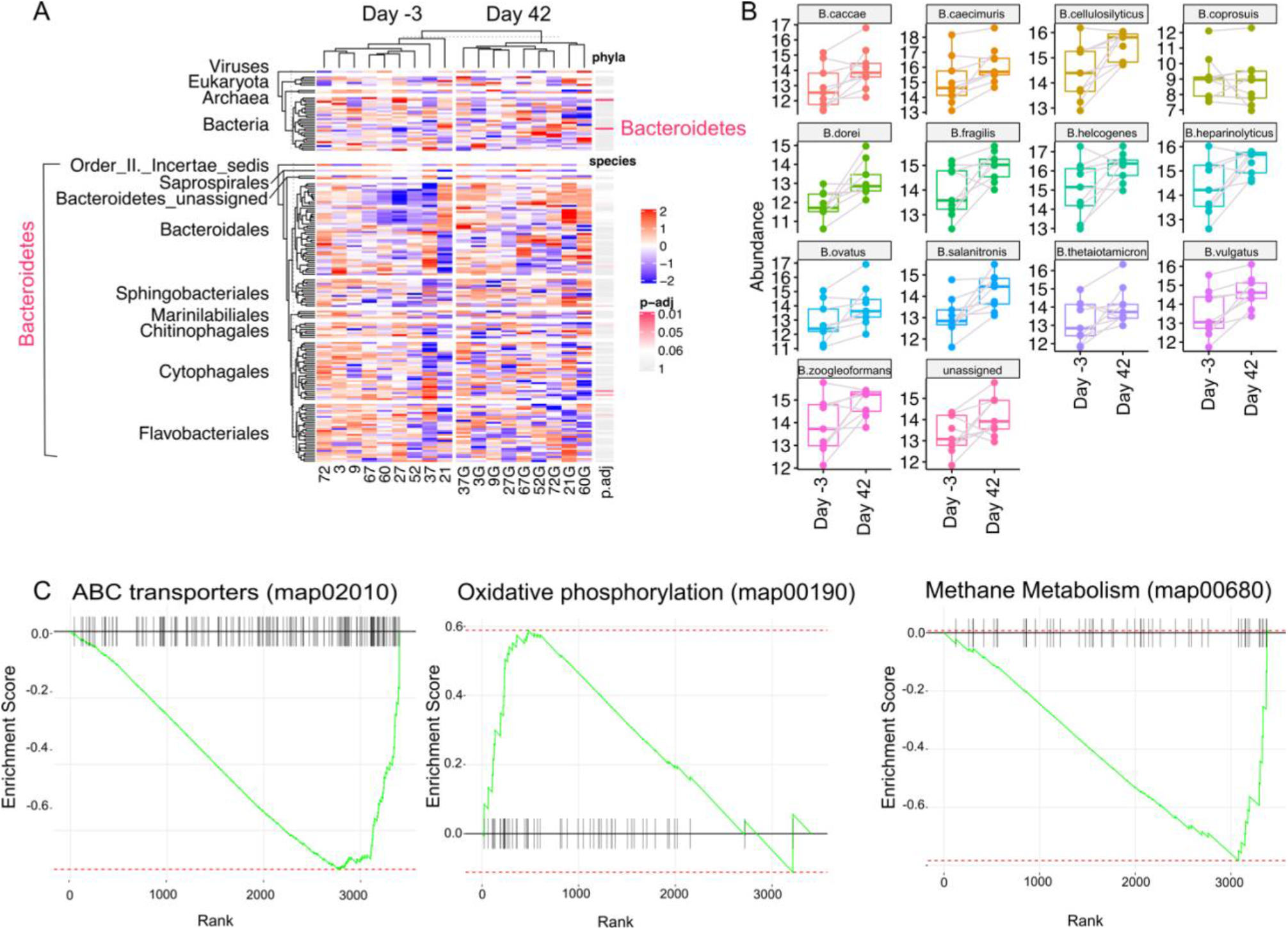
Combination treatment of bupropion and naltrexone impacts on the host gut microbiome. A) Top: Heatmap of Phyla detected in bupropion and naltrexone-treated rats. A pink bar indicates a statistically significant change in abundance by the end of the treatment (p.adj <= 0.05). Notably, abundance of members of the Bacteroidetes phylum were significantly increased at Day 42. To further visualize this increase, all Bacteroidetes members were visualized in the bottom Heatmap, with rows split by Order. Notably, while the Phylum as a whole is significantly increased, only a handful of species are statistically significant (pink bars), but a cluster of Bacteroidales shows a trend towards increasing by the end of the treatment. This heatmap was plotted using ComplexHeatmap, using both phylum and species data [70]. B) Boxplots representing abundance levels of species belonging to the Bacteroides genus indicate a trend towards enrichment by Day 42. Grey lines connect dots corresponding to the same rat. Counts used were log2 transformed and normalized by library size [71]. C) Enrichment plots for the statistically significant pathways “ABC transporters” (map02010; NES = −1.28, p.adj = 0.02), “Oxidative phosphorylation” (map00190; NES = −1.37, p.adj = 0.05) and “Methane metabolism” (map00680; NES = 1.87, p.adj = 0.05). Briefly, genes have been ranked by log2 fold change multiplied by the −log10(p.adjusted), then gsea was performed with the fgsea R/Bioconductor package [72].

**Figure S5.**
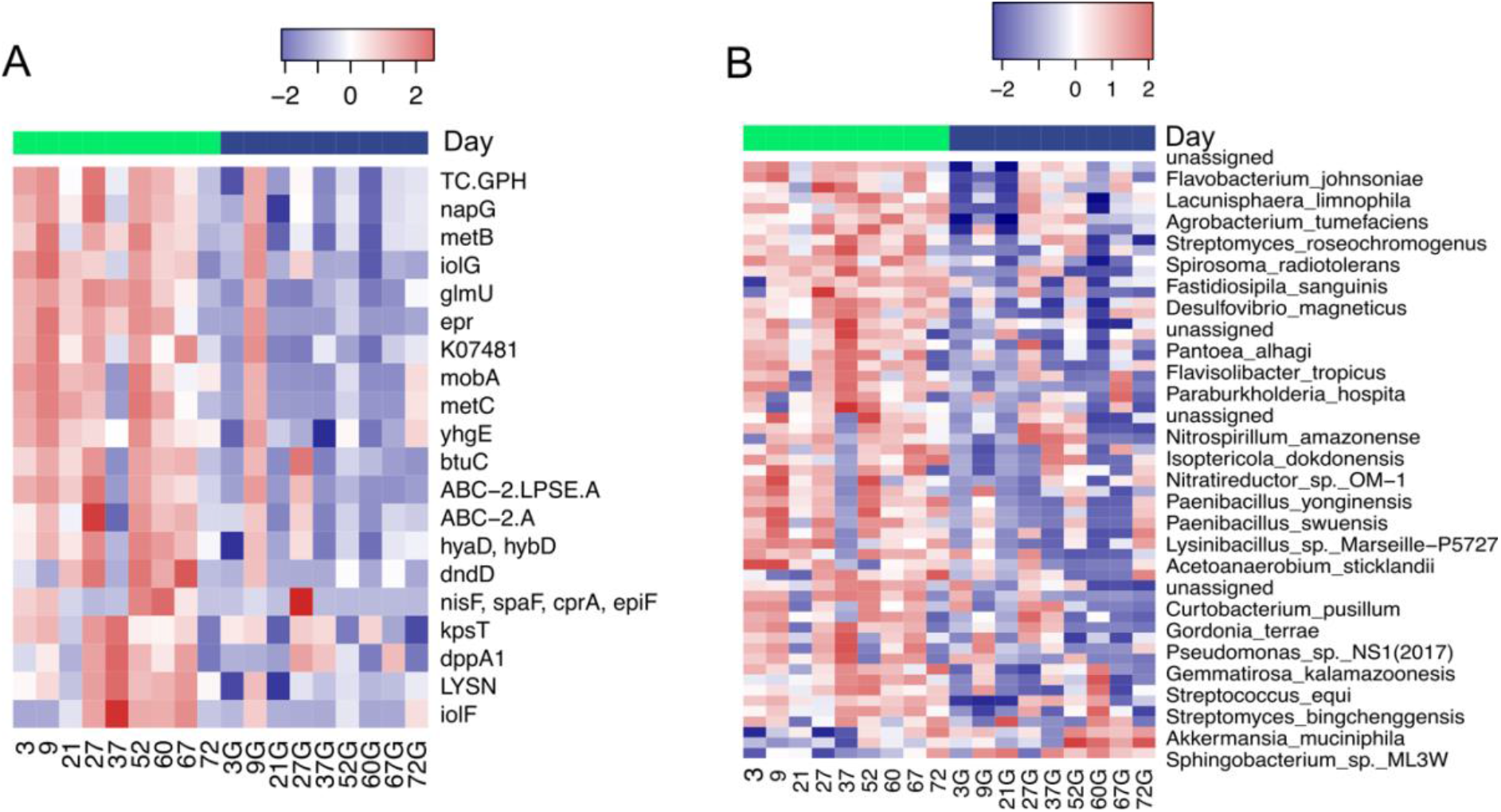
Differentially abundant genes and species upon bupropion and naltrexone combination treatment. A) Differentially abundant genes between bupropion and naltrexone-treated rats at the start and end of the treatment. Heatmap represents log2-transformed and normalized read counts for each of the 20 differentially abundant genes. Significant changes in gene levels were calculated using DESeq2 with the model ~ rat number + Day [73]. P values were corrected with the Benjamini-Hochberg method, and only genes with adjusted p value < = 0. 05 were considered statistically significant. Top bar shows which columns correspond to Day −3 (green) or Day 42 (blue) samples. B) Differentially abundant species between the start and the end of the bupropion and naltrexone combined treatment. Significant changes in species abundance were calculated with DESeq2 with the model ~ rat number + Day [73]. A Heatmap was used to visualize the log2-transformed and normalized read counts for the 57 species which showed an adjusted p.value <= 0.05 and were considered statistically significant. Heatmaps were plotted using the Heatmap3 R/Bioconductor package [41].

**Table S1.**
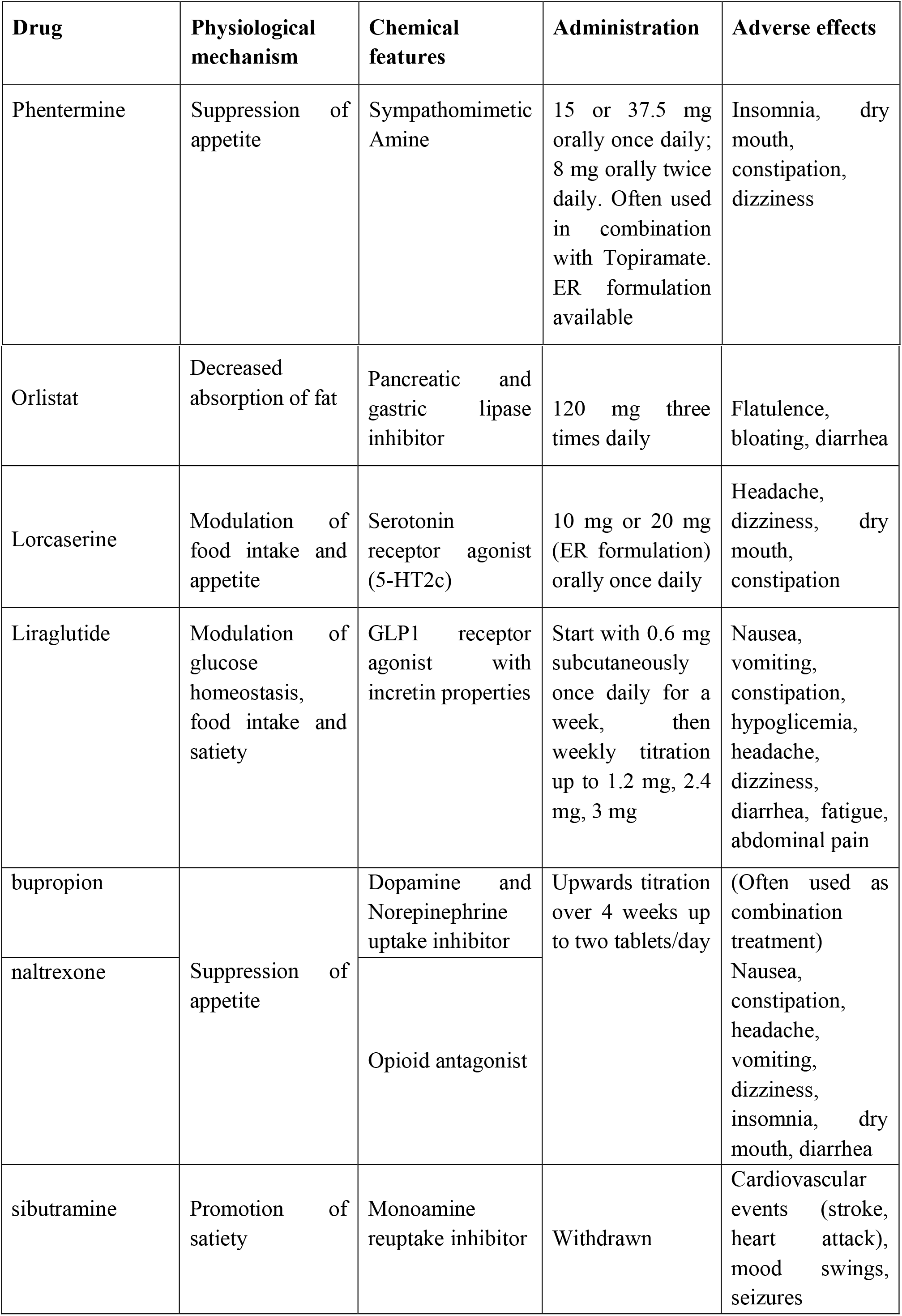
Current pharmacotherapies for obesity. Summary of the most common pharmacological therapies for the treatment of obesity. With the exception of sibutramine, which has been withdrawn from the market due to its cardiological and, to a lesser extent, psychiatric adverse effects, all drugs are currently being used in the clinic, often together with a lifestyle change promoting diet and exercise [6,20].

**Table S2.** Percentage of unassigned reads per sample for representative genes fliC, flgE and motA.

|  | Unassigned reads (%) |  |  |
| --- | --- | --- | --- |
| Sample | *fliC* | *motA* | *flgE* |
| 4 | 53.6082 | 33.3333 | NA |
| 14 | 44.3439 | 100 | 0 |
| 25 | 24.9229 | 13.5647 | 3.44828 |
| 33 | 69.3111 | 41.6452 | 36.2031 |
| 40 | 45.866 | 100 | 47.5207 |
| 55 | 51.7922 | 67.4242 | 85.3261 |
| 62 | 56.2468 | 0 | 7.69231 |
| 75 | 30.7377 | 82.3529 | 52.1866 |
| 85 | 42.9543 | 33.6134 | 38.3754 |
| 4H | 11.8442 | 4.08602 | 0 |
| 14H | 100 | NA | NA |
| 25H | 32.8909 | 3.41686 | 7.20461 |
| 33H | 60.6393 | 51.0417 | 27.6786 |
| 40H | 57.453 | 38.0952 | 32.8125 |
| 55H | 80 | NA | 0 |
| 62H | 6.18557 | 0 | 0 |
| 75H | 30.4888 | 25.7062 | 10.8168 |
| 85H | 63.6459 | 26.875 | 56.5972 |

**Table S3.** Tukey’s post-hoc test significant comparisons following ANOVA on pancreatic insulin levels at Day 42.

| Tukey's post-hoc comparison | Adjusted P.value (Benjamini-Hochberg) |
| --- | --- |
| FK506 vs naltrexone | 0.02 |
| FK506 vs bupropion_naltrexone | 0.02 |
| naltrexone vs FK506_Bupropion | 0.02 |
| naltrexone vs FK506_naltrexone | 0.006 |
| FK506_bupropion vs bupropion_naltrexone | 0.03 |
| FK506_naltrexone vs bupropion_naltrexone | 0.009 |

## Bibliography

1. Wikoff WR, Anfora AT, Liu J, Schultz PG, Lesley SA, Peters EC, et al. Metabolomics analysis reveals large effects of gut microflora on mammalian blood metabolites. Proc Natl Acad Sci U S A. National Academy of Sciences; 2009;106:3698–703.

2. Cryan JF, Dinan TG. Mind-altering microorganisms: The impact of the gut microbiota on brain and behaviour. Nat. Rev. Neurosci. 2012. p. 701–12.

3. Turnbaugh PJ, Hamady M, Yatsunenko T, Cantarel BL, Duncan A, Ley RE, et al. A core gut microbiome in obese and lean twins. Nature. Nature Publishing Group; 2009;457:480–4.

4. Halfvarson J, Brislawn CJ, Lamendella R, Vázquez-Baeza Y, Walters WA, Bramer LM, et al. Dynamics of the human gut microbiome in inflammatory bowel disease. Nat Microbiol [Internet]. Nature Publishing Group; 2017 [cited 2021 Apr 28];2:1–7. Available from: http://www.nature.com/naturemicrobiology

5. WHO. Obesity and overweight [Internet]. [cited 2020 Apr 13]. Available from: https://www.who.int/news-room/fact-sheets/detail/obesity-and-overweight

6. Longo DL, Heymsfield SB, Wadden TA. Mechanisms, Pathophysiology, and Management of Obesity. N Engl J Med New Engl J Med Downloaded n engl j med [Internet]. 2017 [cited 2018 Mar 21];3763:254–66. Available from: http://www.nejm.org/doi/pdf/10.1056/NEJMra1514009

7. Hall JE, Da Silva AA, Do Carmo JM, Dubinion J, Hamza S, Munusamy S, et al. Obesity-induced hypertension: Role of sympathetic nervous system, leptin, and melanocortins [Internet]. J. Biol. Chem. 2010 [cited 2020 Apr 13]. p. 17271–6. Available from: http://www.ncbi.nlm.nih.gov/pubmed/20348094

8. Bornfeldt KE, Tabas I. Insulin resistance, hyperglycemia, and atherosclerosis. Cell Metab. Elsevier; 2011. p. 575–85.

9. Heymsfield SB, Wadden TA. Mechanisms, Pathophysiology, and Management of Obesity. Longo DL, editor. N Engl J Med [Internet]. Massachussetts Medical Society; 2017 [cited 2020 Apr 9];376:254–66. Available from: http://www.nejm.org/doi/10.1056/NEJMra1514009

10. Ley RE, Bäckhed F, Turnbaugh P, Lozupone CA, Knight RD, Gordon JI. Obesity alters gut microbial ecology. Proc Natl Acad Sci U S A. National Academy of Sciences; 2005;102:11070–5.

11. Zhang H, DiBaise JK, Zuccolo A, Kudrna D, Braidotti M, Yu Y, et al. Human gut microbiota in obesity and after gastric bypass. Proc Natl Acad Sci U S A. 2009;106:2365–70.

12. Schwiertz A, Taras D, Schäfer K, Beijer S, Bos NA, Donus C, et al. Microbiota and SCFA in lean and overweight healthy subjects. Obesity. 2010;18:190–5.

13. Nardelli C, Granata I, D’Argenio V, Tramontano S, Compare D, Guarracino MR, et al. Characterization of the Duodenal Mucosal Microbiome in Obese Adult Subjects by 16S rRNA Sequencing. Microorganisms [Internet]. 2020 [cited 2020 Apr 14];8:485. Available from: https://www.mdpi.com/2076-2607/8/4/485

14. Turnbaugh PJ, Ley RE, Mahowald MA, Magrini V, Mardis ER, Gordon JI. An obesity-associated gut microbiome with increased capacity for energy harvest. Nature. Nature Publishing Group; 2006;444:1027–31.

15. Bäckhed F, Ding H, Wang T, Hooper L V., Gou YK, Nagy A, et al. The gut microbiota as an environmental factor that regulates fat storage. Proc Natl Acad Sci U S A. National Academy of Sciences; 2004;101:15718–23.

16. De La Serre CB, Ellis CL, Lee J, Hartman AL, Rutledge JC, Raybould HE. Propensity to high-fat diet-induced obesity in rats is associated with changes in the gut microbiota and gut inflammation. Am J Physiol - Gastrointest Liver Physiol. 2010;299.

17. Liu R, Hong J, Xu X, Feng Q, Zhang D, Gu Y, et al. Gut microbiome and serum metabolome alterations in obesity and after weight-loss intervention. Nat Med [Internet]. Nature Publishing Group; 2017 [cited 2018 Oct 30];23:859–68. Available from: http://www.nature.com/doifinder/10.1038/nm.4358

18. Turnbaugh PJ, Bäckhed F, Fulton L, Gordon JI. Diet-Induced Obesity Is Linked to Marked but Reversible Alterations in the Mouse Distal Gut Microbiome. Cell Host Microbe. 2008;3:213–23.

19. Tremaroli V, Karlsson F, Werling M, Ståhlman M, Kovatcheva-Datchary P, Olbers T, et al. Roux-en-Y Gastric Bypass and Vertical Banded Gastroplasty Induce Long-Term Changes on the Human Gut Microbiome Contributing to Fat Mass Regulation. Cell Metab. 2015;22:228–38.

20. Srivastava G, Apovian CM. Current pharmacotherapy for obesity. Nat Publ Gr [Internet]. 2017 [cited 2018 Mar 21];14. Available from: https://www.nature.com/articles/nrendo.2017.122.pdf

21. Paul Pfluger AT, Kabra DG, Aichler M, De Luca M, Molkentin JD, Tschö Correspondence MH, et al. Calcineurin Links Mitochondrial Elongation with Energy Metabolism. 2015 [cited 2018 Mar 27]; Available from: http://dx.doi.org/10.1016/j.cmet.2015.08.022

22. Ishida H, Mitamura T, Takahashi Y, Hisatomi A, Fukuhara Y, Murato K, et al. Cataract development induced by repeated oral dosing with FK506 (tacrolimus) in adult rats. Toxicology. Elsevier; 1997;123:167–75.

23. Vickers SP, Jackson HC, Cheetham SC. The utility of animal models to evaluate novel anti-obesity agents [Internet]. Br. J. Pharmacol. Br J Pharmacol; 2011 [cited 2021 Feb 25]. p. 1248–62. Available from: https://pubmed.ncbi.nlm.nih.gov/21265828/

24. Greenway FL, Whitehouse MJ, Guttadauria M, Anderson JW, Atkinson RL, Fujioka K, et al. Rational Design of a Combination Medication for the Treatment of Obesity. Obesity [Internet]. John Wiley & Sons, Ltd; 2009 [cited 2021 Feb 25];17:30–9. Available from: http://doi.wiley.com/10.1038/oby.2008.461

25. Greenway FL, Fujioka K, Plodkowski RA, Mudaliar S, Guttadauria M, Erickson J, et al. Effect of naltrexone plus bupropion on weight loss in overweight and obese adults (COR-I): A multicentre, randomised, double-blind, placebo-controlled, phase 3 trial. Lancet [Internet]. 2010 [cited 2020 Oct 26];376:595–605. Available from: http://www.thelancet.com

26. Afgan E, Baker D, Batut B, van den Beek M, Bouvier D, Čech M, et al. The Galaxy platform for accessible, reproducible and collaborative biomedical analyses: 2018 update. Nucleic Acids Res [Internet]. 2018 [cited 2020 Jan 7];46:W537–44. Available from: https://academic.oup.com/nar/article/46/W1/W537/5001157

27. Andrews S. Babraham Bioinformatics - FastQC A Quality Control tool for High Throughput Sequence Data [Internet]. [cited 2020 Jan 7]. Available from: https://www.bioinformatics.babraham.ac.uk/projects/fastqc/

28. Bolger AM, Lohse M, Usadel B. Trimmomatic: a flexible trimmer for Illumina sequence data. Bioinformatics [Internet]. Oxford University Press; 2014 [cited 2018 Jun 12];30:2114–20. Available from: http://www.ncbi.nlm.nih.gov/pubmed/24695404

29. Langmead B, Salzberg SL. Fast gapped-read alignment with Bowtie 2. Nat Methods. 2012;9:357–9.

30. Cribbs AP, Luna-Valero S, George C, Sudbery IM, Berlanga-Taylor AJ, Sansom SN, et al. CGAT-core: a python framework for building scalable, reproducible computational biology workflows. F1000Research [Internet]. F1000 Research Ltd; 2019 [cited 2020 Sep 2];8:377. Available from: https://doi.org/10.12688/f1000research.18674.1

31. Li D, Liu CM, Luo R, Sadakane K, Lam TW. MEGAHIT: An ultra-fast single-node solution for large and complex metagenomics assembly via succinct de Bruijn graph. Bioinformatics [Internet]. Oxford University Press; 2015 [cited 2020 Sep 2];31:1674–6. Available from: https://academic.oup.com/bioinformatics/article/31/10/1674/177884

32. Hyatt D, Chen GL, LoCascio PF, Land ML, Larimer FW, Hauser LJ. Prodigal: Prokaryotic gene recognition and translation initiation site identification. BMC Bioinformatics [Internet]. BioMed Central; 2010 [cited 2020 Sep 2];11:119. Available from: /pmc/articles/PMC2848648/?report=abstract

33. Huerta-Cepas J, Forslund K, Coelho LP, Szklarczyk D, Jensen LJ, Von Mering C, et al. Fast genome-wide functional annotation through orthology assignment by eggNOG-mapper. Mol Biol Evol [Internet]. Oxford University Press; 2017 [cited 2020 Sep 2];34:2115–22. Available from: http://creativecommons.

34. Kanehisa M, Goto S. KEGG: Kyoto Encyclopedia of Genes and Genomes [Internet]. Nucleic Acids Res. Oxford University Press; 2000 [cited 2020 Sep 2]. p. 27–30. Available from: https://pubmed.ncbi.nlm.nih.gov/10592173/

35. Wood DE, Lu J, Langmead B. Improved metagenomic analysis with Kraken 2. Genome Biol [Internet]. BioMed Central Ltd.; 2019 [cited 2020 Sep 2];20:257. Available from: https://genomebiology.biomedcentral.com/articles/10.1186/s13059-019-1891-0

36. Langmead B, Salzberg SL. Fast gapped-read alignment with Bowtie 2. Nat Methods. Nature Publishing Group; 2012;9:357–9.

37. Liao Y, Smyth GK, Shi W. FeatureCounts: An efficient general purpose program for assigning sequence reads to genomic features. Bioinformatics [Internet]. Oxford University Press; 2014 [cited 2020 Sep 2];30:923–30. Available from: https://pubmed.ncbi.nlm.nih.gov/24227677/

38. McMurdie PJ, Holmes S. Phyloseq: An R Package for Reproducible Interactive Analysis and Graphics of Microbiome Census Data. PLoS One. 2013;8.

39. Oksanen J, Kindt R, Legendre P, O’Hara B, Simpson GL, Solymos P, et al. vegan: Community Ecology Package, R package version 2.4-0. R Packag version 22-1 [Internet]. 2016; Available from: http://vegan.r-forge.r-project.org

40. Love MI, Huber W, Anders S. Targeted analysis of nucleotide and copy number variation by exon capture in allotetraploid wheat genome. Genome Biol [Internet]. 2011 [cited 2018 Jun 12];15. Available from: http://www.

41. Zhao S, Yin L, Guo Y, Sheng Q, Shyr Y. Heatmap3: An Improved Heatmap Package. R package version 1.1.7. 2020;

42. Tenenbaum D. KEGGREST: Client-side REST access to KEGG. R package version 1.24.1. 2019;

43. Kassambara A. “ggplot2” Based Publication Ready Plots [R package ggpubr version 0.2.4]. Comprehensive R Archive Network (CRAN);

44. Wickham H, Averick M, Bryan J, Chang W, McGowan L, François R, et al. Welcome to the Tidyverse. J Open Source Softw [Internet]. 2019 [cited 2020 Jan 8];4:1686. Available from: https://joss.theoj.org/papers/10.21105/joss.01686

45. Fisas A, Codony X, Romero G, Dordal A, Giraldo J, Mercé R, et al. Chronic 5-HT6 receptor modulation by E-6837 induces hypophagia and sustained weight loss in diet-induced obese rats. Br J Pharmacol [Internet]. John Wiley and Sons Inc.; 2006 [cited 2021 Feb 25];148:973–83. Available from: /pmc/articles/PMC1751931/

46. De Almeida AR, Monte-Alegre S, Zanini MB, Souza AL, Etchebehere M, Gontijo JAR. Association between prehypertension, metabolic and inflammatory markers, decreased adiponectin and enhanced insulinemia in obese subjects. Nutr Metab [Internet]. BioMed Central Ltd.; 2014 [cited 2020 Dec 8];11:25. Available from: http://nutritionandmetabolism.biomedcentral.com/articles/10.1186/1743-7075-11-25

47. Krzyzanowska K, Zemany L, Krugluger W, Schernthaner GH, Mittermayer F, Schnack C, et al. Serum concentrations of retinol-binding protein 4 in women with and without gestational diabetes. Diabetologia [Internet]. Diabetologia; 2008 [cited 2020 Dec 8];51:1115–22. Available from: https://pubmed.ncbi.nlm.nih.gov/18437353/

48. Rodriguez-Rodriguez AE, Triñanes J, Velazquez-Garcia S, Porrini E, Vega Prieto MJ, Diez Fuentes ML, et al. The Higher Diabetogenic Risk of Tacrolimus Depends on Pre-Existing Insulin Resistance. A Study in Obese and Lean Zucker Rats. Am J Transplant [Internet]. John Wiley & Sons, Ltd; 2013 [cited 2020 Nov 3];13:1665–75. Available from: http://doi.wiley.com/10.1111/ajt.12236

49. Le Chatelier E, Nielsen T, Qin J, Prifti E, Hildebrand F, Falony G, et al. Richness of human gut microbiome correlates with metabolic markers. Nature [Internet]. Nature Publishing Group; 2013 [cited 2020 Oct 30];500:541–6. Available from: https://www.nature.com/articles/nature12506

50. Louis S, Tappu RM, Damms-Machado A, Huson DH, Bischoff SC. Characterization of the gut microbial community of obese patients following a weight-loss intervention using whole metagenome shotgun sequencing. PLoS One. 2016;11:1–18.

51. Subramanian A, Tamayo P, Mootha VK, Mukherjee S, Ebert BL, Gillette MA, et al. Gene set enrichment analysis: A knowledge-based approach for interpreting genome-wide expression profiles. Proc Natl Acad Sci U S A [Internet]. Proc Natl Acad Sci U S A; 2005 [cited 2020 Sep 2];102:15545–50. Available from: https://pubmed.ncbi.nlm.nih.gov/16199517/

52. Neville BA, Sheridan PO, Harris HMB, Coughlan S, Flint HJ, Duncan SH, et al. Pro-Inflammatory Flagellin Proteins of Prevalent Motile Commensal Bacteria Are Variably Abundant in the Intestinal Microbiome of Elderly Humans [Internet]. PLoS One. Public Library of Science; 2013 [cited 2020 Sep 2]. Available from: /pmc/articles/PMC3720852/?report=abstract

53. Hou YP, He QQ, Ouyang HM, Peng HS, Wang Q, Li J, et al. Human Gut Microbiota Associated with Obesity in Chinese Children and Adolescents. Biomed Res Int. Hindawi Limited; 2017;2017.

54. Minamino T, Morimoto Y V., Kawamoto A, Terashima H, Imada K. *Salmonella* Flagellum. Salmonella - A Re-emerging Pathog [Internet]. InTech; 2018 [cited 2020 Sep 2]. Available from: http://dx.doi.org/10.5772/intechopen.73277

55. Bergeron JR. Structural modeling of the flagellum MS ring protein FliF reveals similarities to the type III secretion system and sporulation complex. PeerJ [Internet]. PeerJ Inc.; 2016 [cited 2020 Sep 2];2016. Available from: /pmc/articles/PMC4768692/?report=abstract

56. Takekawa N, Terahara N, Kato T, Gohara M, Mayanagi K, Hijikata A, et al. The tetrameric MotA complex as the core of the flagellar motor stator from hyperthermophilic bacterium. Sci Rep [Internet]. Nature Publishing Group; 2016 [cited 2020 Sep 2];6:1–8. Available from: http://www.nature.com/scientificreports

57. Barker CS, Meshcheryakova I V., Kostyukova AS, Samatey FA. FliO Regulation of FliP in the Formation of the Salmonella enterica Flagellum. Dutcher SK, editor. PLoS Genet [Internet]. Public Library of Science; 2010 [cited 2020 Sep 2];6:e1001143. Available from: https://dx.plos.org/10.1371/journal.pgen.1001143

58. Mikami A, Ogita T, Namai F, Shigemori S, Sato T, Shimosato T. Oral administration of Flavonifractor plautii attenuates inflammatory responses in obese adipose tissue. Mol Biol Rep [Internet]. Springer; 2020 [cited 2020 Sep 2];1:3. Available from: https://doi.org/10.1007/s11033-020-05727-6

59. Castaner O, Goday A, Park YM, Lee SH, Magkos F, Shiow SATE, et al. The gut microbiome profile in obesity: A systematic review. Int J Endocrinol [Internet]. Hindawi Limited; 2018 [cited 2020 Sep 2];2018. Available from: /pmc/articles/PMC5933040/?report=abstract

60. Peters BA, Shapiro JA, Church TR, Miller G, Trinh-Shevrin C, Yuen E, et al. A taxonomic signature of obesity in a large study of American adults. Sci Rep. Nature Publishing Group; 2018;8:1–13.

61. Lai ZL, Tseng CH, Ho HJ, Cheung CKY, Lin JY, Chen YJ, et al. Fecal microbiota transplantation confers beneficial metabolic effects of diet and exercise on diet-induced obese mice. Sci Rep [Internet]. Nature Publishing Group; 2018 [cited 2020 Sep 3];8:14. Available from: http://www.nature.com/scientificreports

62. Liu R, Hong J, Xu X, Feng Q, Zhang D, Gu Y, et al. Gut microbiome and serum metabolome alterations in obesity and after weight-loss intervention. Nat Med. 2017;23:859–68.

63. Kootte RS, Levin E, Salojärvi J, Smits LP, Hartstra A V., Udayappan SD, et al. Improvement of Insulin Sensitivity after Lean Donor Feces in Metabolic Syndrome Is Driven by Baseline Intestinal Microbiota Composition. Cell Metab. Cell Press; 2017;26:611–619.e6.

64. Gershon MD, Tack J. The Serotonin Signaling System: From Basic Understanding To Drug Development for Functional GI Disorders. Gastroenterology [Internet]. W.B. Saunders; 2007 [cited 2020 Dec 8];132:397–414. Available from: https://pubmed.ncbi.nlm.nih.gov/17241888/

65. Yanovski SZ, Yanovski JA. Obesity. Wood AJJ, editor. N Engl J Med [Internet]. Massachusetts Medical Society; 2002 [cited 2020 Dec 8];346:591–602. Available from: http://www.nejm.org/doi/10.1056/NEJMra012586

66. Yano JM, Yu K, Donaldson GP, Shastri GG, Ann P, Ma L, et al. Indigenous bacteria from the gut microbiota regulate host serotonin biosynthesis. Cell. Cell Press; 2015;161:264–76.

67. Carvalho FA, Koren O, Goodrich JK, Johansson MEV, Nalbantoglu I, Aitken JD, et al. Transient inability to manage proteobacteria promotes chronic gut inflammation in TLR5-deficient mice. Cell Host Microbe [Internet]. Cell Host Microbe; 2012 [cited 2020 Sep 4];12:139–52. Available from: https://pubmed.ncbi.nlm.nih.gov/22863420/

68. Vijay-Kumar M, Aitken JD, Carvalho FA, Cullender TC, Mwangi S, Srinivasan S, et al. Metabolie syndrome and altered gut microbiota in mice lacking toll-like receptor 5. Science (80-) [Internet]. Science; 2010 [cited 2020 Sep 4];328:228–31. Available from: https://pubmed.ncbi.nlm.nih.gov/20203013/

69. Dalby MJ, Ross AW, Walker AW, Morgan PJ. Dietary Uncoupling of Gut Microbiota and Energy Harvesting from Obesity and Glucose Tolerance in Mice. Cell Rep. Elsevier B.V.; 2017;21:1521–33.

70. Gu Z, Eils R, Schlesner M. Complex heatmaps reveal patterns and correlations in multidimensional genomic data. Bioinformatics. Oxford University Press; 2016;32:2847–9.

71. Love MI, Huber W, Anders S. Moderated estimation of fold change and dispersion for RNA-seq data with DESeq2. Genome Biol [Internet]. BioMed Central Ltd.; 2014 [cited 2020 Jun 16];15:550. Available from: http://genomebiology.biomedcentral.com/articles/10.1186/s13059-014-0550-8

72. Sergushichev AA. An algorithm for fast preranked gene set enrichment analysis using cumulative statistic calculation. bioRxiv [Internet]. Cold Spring Harbor Laboratory; 2016 [cited 2020 Jun 17];060012. Available from: http://dx.doi.org/10.1101/060012 https://www.biorxiv.org/content/10.1101/060012v1

73. Love MI, Huber W, Anders S. Moderated estimation of fold change and dispersion for RNA-seq data with DESeq2. Genome Biol [Internet]. 2014 [cited 2018 Mar 28];15. Available from: http://www.

74. Kassambara A. rstatix: Pipe-Friendly Framework for Basic Statistical Tests. R package version 0.6.0. 2020.

